# Co-occurrence of diverse defense systems shapes complex microbe-virus relationships in deep-sea cold seeps

**DOI:** 10.1101/2024.09.26.614923

**Authors:** Yingchun Han, Jing Liao, Chengpeng Li, Fengmin Xing, Jiaxue Peng, Xinyue Liu, Wentao Xie, Fabai Wu, Huahua Jian, Rui Cheng, Xiyang Dong

**Author notes:** Correspondence can be addressed to Xiyang Dong and Rui Cheng. These authors contributed equally to this work (Yingchun Han, Jing Liao).

## Abstract

Cold seeps host diverse microbes and viruses with numerous unexplored defense and anti-defense systems. Analysis of 3,813 microbial and 13,336 viral genomes from 191 metagenomes across 17 cold seep sites reveals extensive microbial defense repertoires, with over 60% representing candidate systems. Experimental validation confirms that several candidates protect against viral infection. These defense systems frequently co-occur, suggesting potential synergistic interactions, and are broadly distributed across sediments. In response, viruses have evolved diverse anti-defense genes, and the concurrent presence of multiple viral and microbial systems highlights intricate coevolution. Functionally critical lineages, such as anaerobic methanotrophic archaea, sulfate-reducing bacteria, and diazotrophs, appear to modify their defensive strategies under ecological and environmental pressures; for example, sulfate-reducing bacteria harbor multiple Gabija systems while corresponding viruses carry anti-Gabija genes, illustrating specific coevolutionary adaptations. Overall, these findings underscore the critical role of virus-microbe interactions in shaping microbial metabolic functions and environmental adaptation in deep-sea ecosystems.

## Introduction

Cold seeps are extreme deep-sea environments located along continental margins worldwide, characterized by the migration of fluids rich in methane and other hydrocarbons from the deep subsurface to the sediment-seawater interface^1, 2^. The continuous release of these fluids enhances local biodiversity and microbial activity, making cold seeps thriving deep-sea oases^2^. These ecosystems support diverse microbial communities, including anaerobic methanotrophic archaea (ANME) and their syntrophic sulfate-reducing bacterial partners^3, 4^. Moreover, the sediments harbor a rich diversity of previously understudied prokaryotes and viruses^5, 6^, suggesting the existence of complex and novel microbe-virus interactions within cold seep ecosystems. Prokaryotes in these environments constantly face viral threats, leading to the evolution of multiple defense mechanisms that regulate the flow of genetic information spread by mobile genetic elements (MGEs) through horizontal gene transfer (HGT)^7–9^. Cold seep viruses also employ diverse strategies for environmental adaptation, including counter-defense systems, auxiliary metabolic genes (AMGs), reverse transcriptases, and alternative genetic code assignments^10^. Among these, anti-defense systems have evolved to counteract the defense mechanisms of host microbes. Despite these insights, the diversity of prokaryotic defense genes and the intricate evolutionary interactions between prokaryotes and viruses in cold seep ecosystems remain largely unexplored.

Prokaryotes have developed an arsenal of defense systems to provide partial or full resistance against viral and MGE infections^7, 8, 11^. The most widespread prokaryotic immune systems are the restriction-modification (RM) systems and the clustered regularly interspaced short palindromic repeats CRISPR-Cas system. RM systems recognize specific DNA sequences in invading phages and distinguish self from non-self DNA through methylation^12^, while CRISPR-Cas provides acquired immunity by incorporating short viral and mobile genetic element (MGE) sequences as spacers in the host genome^9, 13^. In recent years, many novel prokaryotic antiviral systems have been revealed. For example, Gabija system, composed of GajA and GajB proteins, degrades viral and host DNA through site-specific double-stranded DNA nicking by GajA upon activation^14^. The cyclic oligonucleotide-based antiphage signaling system (CBASS) uses cyclic oligonucleotides to activate antiviral immunity^15^. Gabija and CBASS systems defend against phages by inducing cell death or metabolic disturbance upon infection, an abortive infection mechanism that is similarly adopted by various other defense systems^14, 16^. Recent efforts have uncovered several defense systems with diverse genetic architectures^7, 8^, reflecting the strong selective pressure imposed by viruses and MGEs on microbial communities in various ecosystems such as soil, marine, human gut and rumen^9, 17^. However, in the deep sea, particularly in cold seep environments, there is a lack of corresponding studies on microbial defense systems, despite these environments potentially equipped with vast and untapped wealth of immune systems and mechanisms^5, 6^.

Defense genes encoding components from multiple distinct systems tend to cluster together in specific genomic regions referred to as defense islands^9, 18, 19^. This co-localization, especially within MGEs, facilitates joint HGT, which is thought to confer fitness advantages to recipient microbes, particularly in virus-rich environments^20^. Synergistic interactions between different defense systems may drive their co-localization and favor their joint transfer^21, 22^, as evidenced by the conservation of certain sets of defense systems across different microbial species. Synergistic systems enhance protective effects against specific phages beyond the sum of individual systems, providing microbes with an evolutionary advantage. Recent research indicated that non-random synergistic interactions between defense systems in *Escherichia coli* are adaptive responses to specific selective forces from the virome, shaping unique fitness landscapes in their specific ecological niches^23^. Given that cold seep viruses in extreme deep-sea subseafloor conditions are endemic and unique compared to those in other ecosystems^6, 10^, the specific virus-driven synergistic interactions between defense systems may differ and warrant investigation.

Viruses can evade microbial defenses both indirectly, by mutating immune system activators or disrupting defense pathways, and directly, by deploying anti-defense genes that target immune system components^24^. In this ongoing arms race, viruses have evolved sophisticated anti-defense mechanisms to overcome microbial defense systems^24, 25^, such as anti-CRISPRs (Acr)^26, 27^, anti-RM^28^, anti-CBASS^29^ and anti-Gabija^30^. These viral anti-defense systems not only highlight the complexity of viral adaptations but also reveal evolutionary connections within microbial immunity^24^. Beyond defense evasion, viruses play a crucial role in shaping microbial communities and influencing ecosystem functioning, including key biogeochemical cycles such as methane oxidation, sulfate reduction and nitrogen fixation, often mediated through AMGs^31, 32^. Despite their importance, the interactions between viral anti-defense mechanisms and functionally critical lineages in the cold seep environment^25^—with its darkness, low temperatures, and hydrocarbon-rich conditions—is not well understood. Understanding these dynamics in extreme deep-sea ecosystem is vital for unraveling the full impact of viruses on microbial ecology and biogeochemical processes.

To explore the diversity of cold seep microbial defense systems, their interactions, and the evolutionary associations between these systems and viral anti-defense strategies in extreme deep-sea subseafloor habitats, we analyze 3,813 microbial genomes and 13,336 viral genomes from 191 metagenomic samples collected at 17 global sites **(Supplementary Fig. 1)**. Our findings reveal that cold seep microbes possess a vast and diverse repertoire of defense systems, which may work synergistically against viral infections. In response, cold seep viruses deploy diverse anti-defense systems to counter microbial defenses. This complex virus-microbe interaction drives microbial evolution and enhances environmental adaptation. Overall, this study enriches our understanding of the ecological and evolutionary dynamics of virus-microbe interactions in deep sea cold seep subsurface ecosystems.

## Results and Discussion

### Cold seep prokaryotes possess diverse and putative defense systems

We investigated defense strategies of cold seep microbiome by analyzing 3,813 species-level metagenome-assembled genomes (MAGs), comprising 639 archaeal and 3,174 bacterial genomes, spanning 107 bacterial and 16 archaeal phyla **(Supplementary Fig. 2)**. The bacterial phyla with the largest diversity of recovered species included *Pseudomonadota* (n = 471), *Chloroflexota* (n = 399), *Desulfobacterota* (n = 298), *Planctomycetota* (n = 225), and *Bacteroidota* (n = 219). Among the archaeal phyla, *Halobacteriota* (n = 153), *Thermoplasmatota* (n = 119), *Thermoproteota* (n = 110), and *Asgardarchaeota* (n = 105) were the most diverse.

We identified defense genes and systems using DefenseFinder^19^ and PADLOC^33^ and detected a total of 36,783 defense genes in 65% of cold seep microbial genomes, which were assigned to 26,389 defense systems **(Fig. 1a and Supplementary Data 1-3)**. Among these, 63% (n = 16,520) were annotated as candidate systems, primarily including HECs (Hma-Embedded Candidates) and PDCs (Phage Defense Candidates). These candidates were predicted by PADLOC based on their recurrent genetic embedding within known defense loci (e.g., Hma, RM, BREX, DISARM) rather than defense-island enrichment^34^. Together, these candidates comprised 18,195 defense genes, highlighting that cold seep microbes harbor a diverse array of putative antiviral strategies. Experimentally verified defense systems, comprising 37% (n = 9,869), included 106 types, with RM systems (n = 3,778) being the most prevalent, followed by CRISPR-Cas (n = 977), SoFic (n = 715), and AbiE (n = 612). This distribution aligns with previous findings that RM and CRISPR-Cas systems are predominant in microbiomes from soil, human gut, and Haima cold seep environments^9, 10^. In contrast, seawater microbial genomes, while also rich in RM systems, exhibit significant enrichment of CBASS^9^, suggesting distinct distribution patterns of defense systems in cold seep sediments compared to other marine environments.

**Figure 1.**
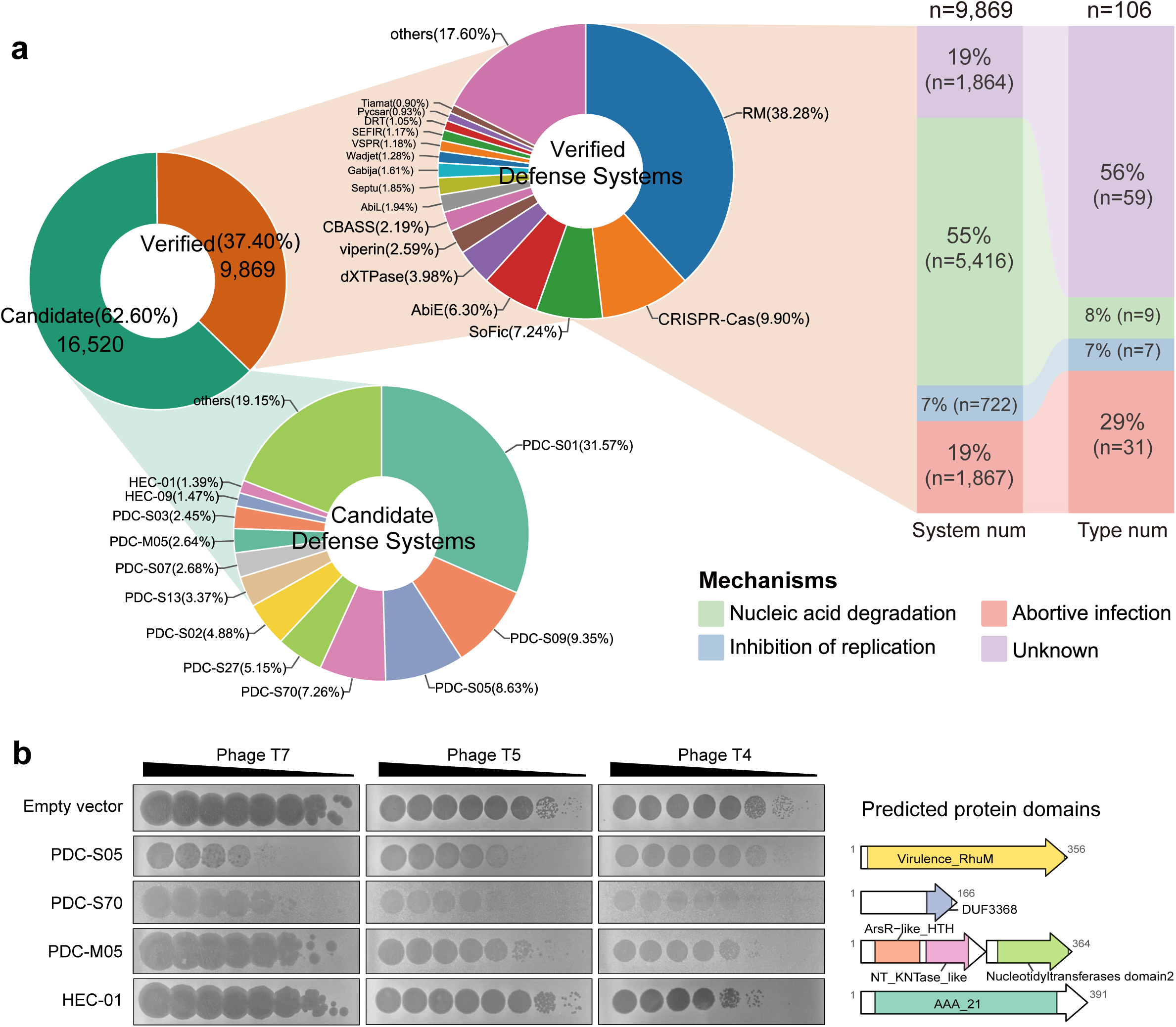
Distribution and activities of defense systems in cold seep prokaryotes. **(a)** Types and proportions of verified defense systems and candidate defense systems in cold seep microbial genomes, including their primary mechanisms of action and corresponding system counts. “DS” denotes defense systems and “num” represents the total count. “Type num” refers to the number of distinct defense system types within each mechanism category. **(b)** Plaque assays demonstrating phage infection of *Escherichia coli* B (ATCC^®^ 11303™) transformed with plasmids carrying various candidate defense systems or an empty vector (control) using a small volume drop method. The bacteria described above were mixed in the molten agar matrix and ten-fold serial dilutions of phages T7, T5, and T4 were spotted onto bacterial lawns. All the assays are repeated at least 3 times with similar results. Predicted protein domains of the tested systems are shown on the right. Detailed data on annotated defense systems are provided in **Supplementary Data 1-3**.

To validate whether these candidate systems confer antiviral activity, we randomly selected eight candidate systems from the top 12 most abundant system types (each accounting for more than 1% of the total), including PDC-S01, PDC-S09, PDC-S05, PDC-S70, PDC-S27, PDC-S02, PDC-M05, and HEC-01. Within each selected type, one representative gene cluster was randomly chosen for experimental validation. Recombinant vectors containing these candidate systems were transformed into the host strain *E. coli* B (ATCC^®^ 11303™), which naturally lacks these systems. The transformed strains were then challenged with three representative lytic phages: T7 (*Podoviridae*), T5 (*Siphoviridae*), and T4 (*Myoviridae*). Among them, four candidate defense systems (PDC-S05, PDC-S70, PDC-M05, and HEC-01) conferred protection against at least one phage **(Fig. 1b)**. Especially, the PDC-S05 and PDC-S70 systems exhibited obvious and broad-spectrum phage resistance. The PDC-S05 system showed an approximately 1,000-fold reduction in efficiency of plating (EOP) for phages T7 and T5, and a ∼10-fold decrease for phage T4. Similarly, the PDC-S70 system caused a ∼100-fold decrease in EOP for phages T7 and T4, and an ∼1,000-fold decrease for phage T5. Although these results confirmed the antiviral activity of four systems, the vast, yet candidate defense systems require further experimental validation. Predicted protein domain analysis of these four systems **(Supplementary Data 4)** revealed that HEC-01 contains an AAA_21 domain, which is associated with ATP binding and hydrolysis activity. PDC-M05A harbors ArsR-like_HTH and NT_KNTase_like domains, suggesting DNA-binding transcription factor activity. Additionally, PDC-S05 features a Virulence_RhuM domain, which is linked to virulence and pathogenicity **(Fig. 1b)**. These findings indicate that most of these candidate defense systems may function through mechanisms such as nucleic acid degradation and replication inhibition.

Defense system mechanisms can be categorized in three types based on their fundamental mechanisms of action: nucleic acid degradation, abortive infection, inhibition of replication, and other unknown mechanisms^19, 35^. For instance, RM and CRISPR-Cas systems function through viral nucleic acid degradation, CBASS and retrons through abortive infection, and Viperins and dCTP deaminases through inhibition of replication. Among the experimentally confirmed defense systems in cold seep microbes **(Fig. 1a)**, the most prevalent mechanism was nucleic acid degradation (n = 5,416, 55%), followed by abortive infection (n = 1,867, 19%) and replication inhibition (n = 722, 7%). Notably, a substantial proportion of defense mechanisms remain unidentified (n = 1,864, 19%), encompassing 59 types that represent 56% of all detected defense system types (n = 106).

### Cold seep defensive patterns are associated with microbial taxonomy

The distribution of verified defense systems differed across taxonomical domains **(Fig. 2a and Supplementary Fig. 3)**, suggesting that archaea and bacteria possess diverse and distinct defense strategies against viral or MGE infections^20, 33, 36^. Most bacterial and archaeal MAGs harbored a limited number of defense systems, averaging four per genome **(Fig. 2b and 2c)**, consistent with observations from other marine environments^9, 10^. However, this value is lower than the average numbers reported for bacterial genomes (5.6-5.8)^8, 19^ and archaeal genomes (4.55)^37^. This discrepancy may be attributed to the completeness of MAGs used in this study, differences in detection methods and parameter settings, and the unique ecological pressures of cold seep environments. On average, bacterial genomes in cold seep environments still encode more defense systems per genome than archaeal genomes (4.1 vs 3.5, **Supplementary Data 5**). Despite this average, several genomes (n = 40) encoded more than 20 defense systems. For instance, a genome belonging to the *Bacteroidales* order encodes 54 defense systems, followed by 48 in a *Methylococcales* genome (*Gammaproteobacteria*) and 39 in a *Desulfuromonadales* genome (*Desulfobacterota*) **(Supplementary Data 3)**. Additionally, we found that the majority of defense systems (∼99.88%) are encoded in chromosomes, several systems—especially AbiV, AbiD, DRT, and AbiL—are carried by plasmids, and others like AbiZ, PT_Dnd, and Lamassu are carried by proviruses **(Fig. 3 and Supplementary Fig. 4a)**. Furthermore, the number of defense systems in prokaryotes increases with the number of proviruses and plasmids, a trend particularly evident in bacteria **(Supplementary Fig. 4b)**, suggesting that defense genes in cold seep prokaryotes could be disseminated between microbes through MGEs^38, 39^.

**Figure 2.**
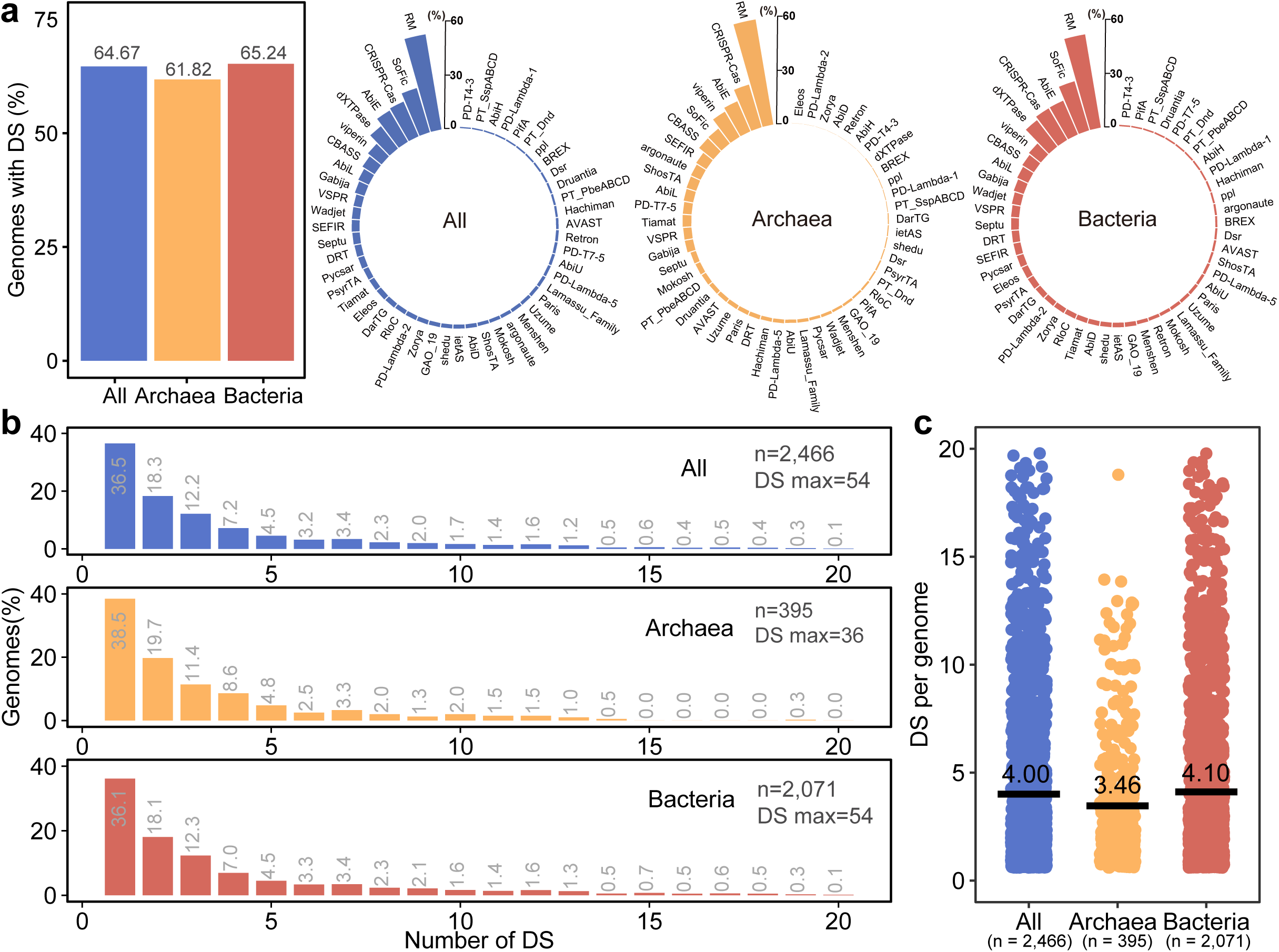
Characteristics of defense systems in cold seep prokaryotes. **(a)** Proportion of metagenome-assembled genomes (MAGs) containing defense systems among cold seep prokaryotes, and the proportions of various types of defense systems within all prokaryotes, archaeal, and bacterial MAGs. **(b)** Distribution of the number of defense systems (DS) per MAG for all prokaryotes (blue), archaea (orange), and bacteria (red). **(c)** Number of defense systems (DSs) per MAG for all prokaryotes (blue, n = 2,466), archaea (orange, n = 395), and bacteria (red, n = 2,071), with black lines indicating the average number. Detailed statistical data are provided in **Supplementary Data 5**. Detailed data on the proportion of genomes carrying defense systems are provide as a Source Data file.

**Figure 3.**
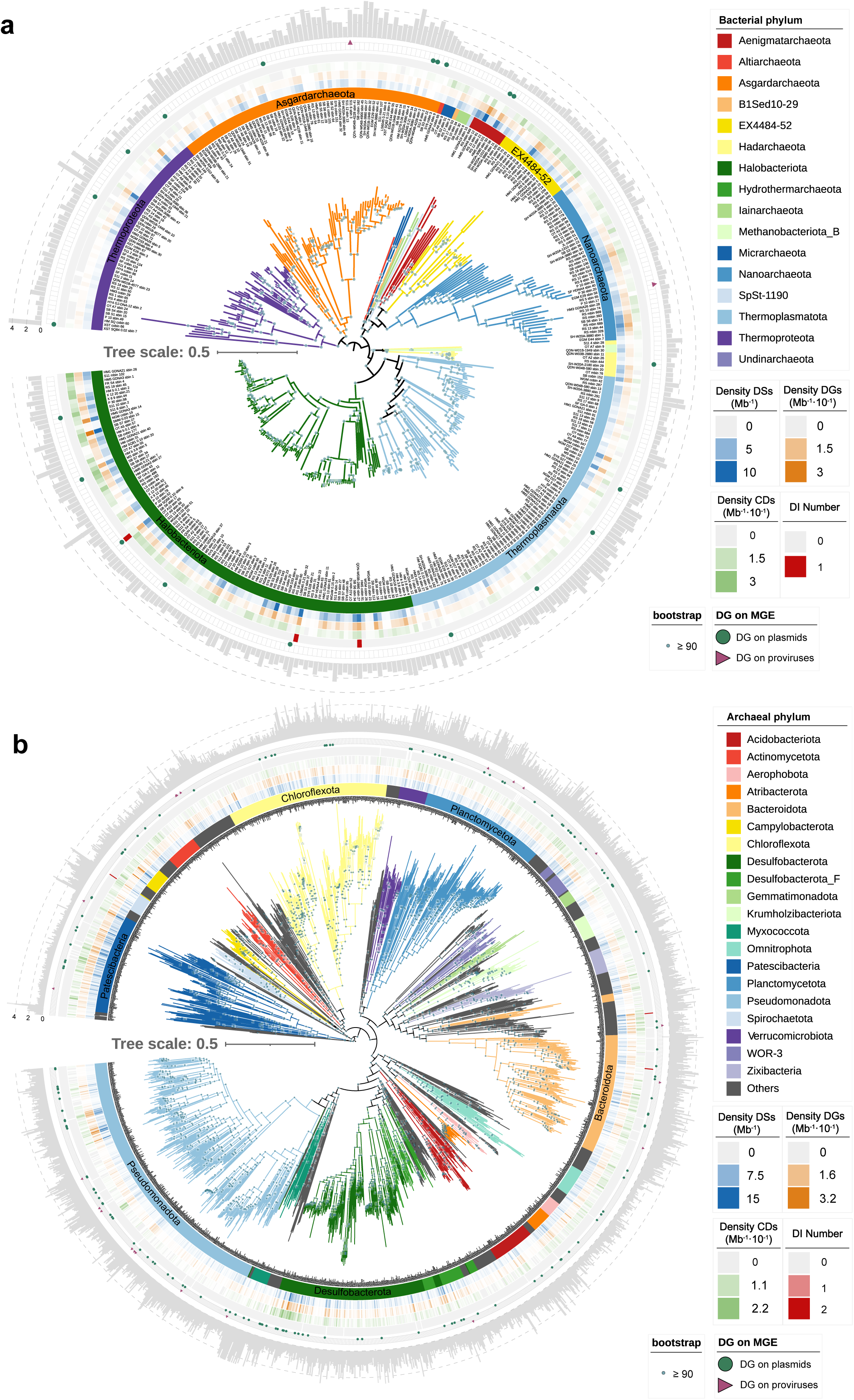
Abundance and distribution of defense systems in cold seep prokaryotes. Phylogenetic representation of archaeal **(a)** and bacterial **(b)** MAGs, categorized and color-coded by their corresponding phyla. The heatmap illustrates the density of defense systems (DSs per Mb, blue), defense genes (DGs per Mb×10^-1^, orange), candidate defense systems (CDs per Mb×10^-1^, green), and the number of defense islands (DIs, red). Circles and triangles denote MAGs with defense systems located on plasmids and proviruses, respectively. The outer bar charts display the genome sizes of MAGs (Mb). Detailed statistical data for this figure are provided in **Supplementary Data 3**.

The density of defense systems (per MAG and per Mb) varied widely among different clades **(Fig. 3, Supplementary Figs. 5-6, and Supplementary Data 3)**. In archaeal MAGs, the density ranged from as low as 0.11 in certain phyla to over 10.5 in the *Aenigmatarchaeota* phylum (**Fig. 3a and Supplementary Fig. 5**). In bacterial MAGs, the density can be as high as 15.1 in the *Desulfobacterota* phylum (**Fig. 3b and Supplementary Fig. 6**). While previous studies have suggested that deeper phylogenetic lineages accumulate more horizontally transferred genes, potentially leading to a buildup of defense systems^23^, our linear regression analyses indicate that phylogenetic depth has a relatively minor impact on the number of defense systems **(Supplementary Data 6)**. This suggests that other factors, such as environmental pressures, viral diversity, and ecological interactions, may play a more significant role in shaping the abundance and distribution of defense systems.

### Cold seep defense systems tend to co-occur and may interact synergistically

Previous studies have demonstrated that certain defense systems interact to enhance or broaden protection against viruses^23, 40, 41^. Here, based on the relative abundances of defense systems, we used Pearson correlation analysis to investigate the relationships among defense systems in cold seep prokaryotes to explore their potential interactions. We considered only statistically significant correlations (*p* < 0.05), following the approach of Wu et al.^42^. We identified 182 pairs of defense systems exhibiting significant positive correlations and 10 pairs with significant negative correlations **(Fig. 4a and Supplementary Data 7**). Among these significantly correlated pairs, 94.8% were positive, suggesting that defense systems are more likely to co-occur rather than to be negatively associated. This pattern is consistent with observations in certain bacterial genomes, such as *E. coli*, *Enterobacterales, Bacillales, Burkholderiales,* and *Pseudomonadales*^23^. Notably, prevalent defense systems such as Gabija (20 pairs), RM (19 pairs), CRISPR-Cas (19 pairs), and SoFic (18 pairs) exhibited a higher number of significant co-occurrences. Additionally, less common systems such as Septu (18 pairs), Druantia (15 pairs), and Menshen (15 pairs), also demonstrated a tendency to co-occur with other defense systems. This suggests that the significant co-occurrence of these systems is not merely a result of their higher overall frequency but may also stem from functional complementarities^22, 41^. For instance, CRISPR-Cas and RM systems can work synergistically by targeting different sites on invading phage DNA, thereby enhancing overall protection^41^. Similarly, defense islands encoding both BREX systems and type IV restriction enzymes have been shown to provide layered protection^22^. Furthermore, RM-mediated cleavage of viral DNA can stimulate spacer acquisition in CRISPR-Cas systems, reinforcing their cooperative function^43^. Interestingly, different subtypes of the same defense system exhibited distinct co-occurrence patterns with other systems **(Supplementary Fig. 7 and Supplementary Data 8)**. For example, Wadjet I co-occurred with multiple other defense systems, showing 11 pairs of co-occurrences. In contrast, Wadjet II had only 4 co-occurrences, which included CRISPR-Cas I-G, CBASS III, Eleos, and PD-T4-3; notably, none of these systems co-occurred with Wadjet I. These specific co-occurrence patterns may indicate unique cooperative interactions among different subtypes of defense systems.

**Figure 4.**
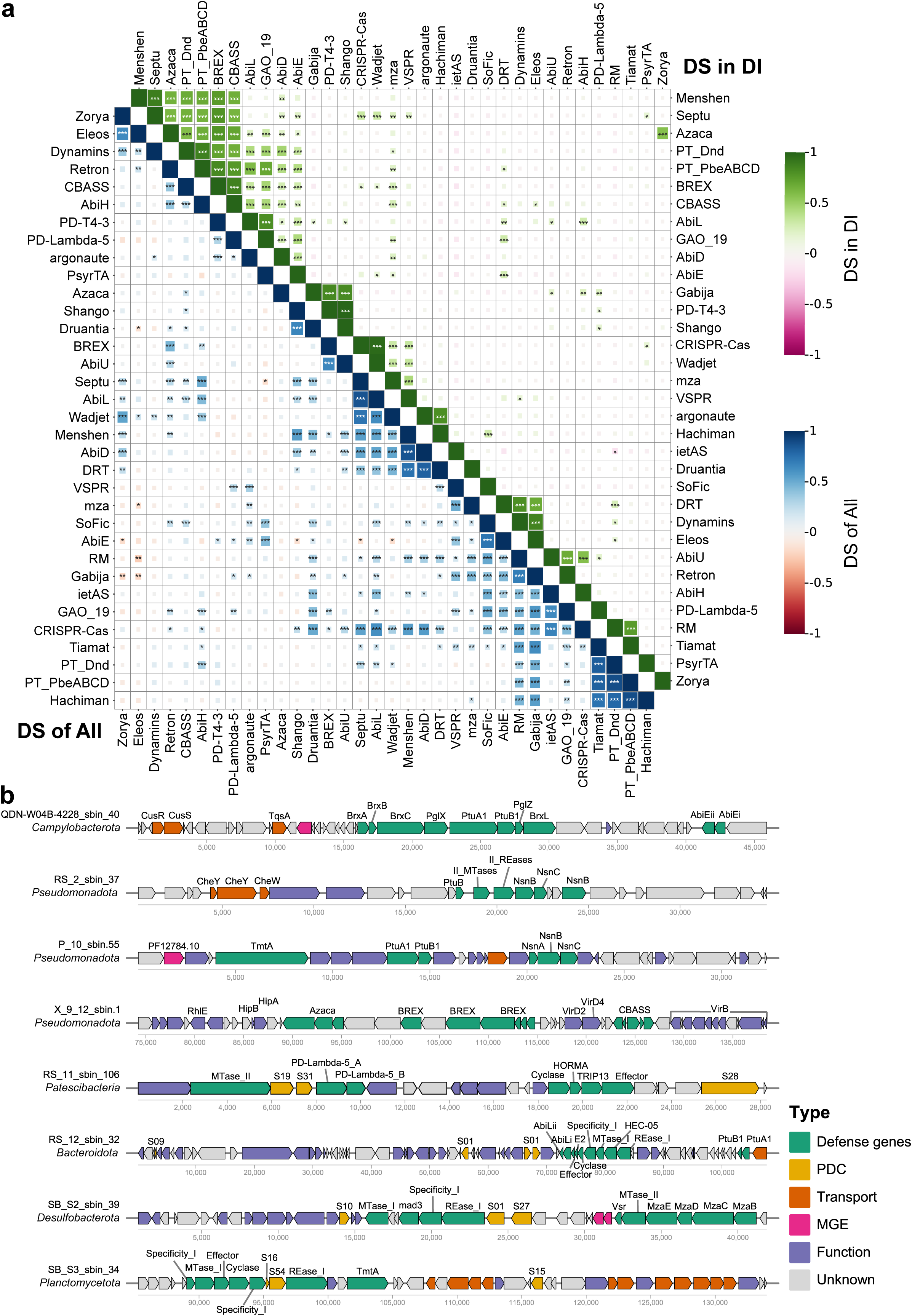
Co-occurrence patterns among defense systems in cold seep prokaryotes. **(a)** Correlations among 34 defense systems in cold seep prokaryotes. The lower-left matrix (blue-red gradient) displays correlations among all defense systems across prokaryotic MAGs, while the upper-right matrix (green-pink gradient) shows correlations within defense islands. Pearson correlation coefficients were calculated based on system abundance using two-sided tests. Coefficients are color-coded, and larger squares indicate stronger correlations. Significance levels are denoted by asterisks: * *p* ≤ 0.05, ** *p* ≤ 0.01, *** *p* ≤ 0.001. **(b)** Genomic collinearity of eight contiguous fragments containing defense islands. Defense genes are highlighted in green, candidate defense genes in yellow, and associated genes are color-coded: transport-related genes (orange), mobile genetic elements (MGEs; pink), other annotated functional genes (purple), and unknown functions (gray). Detailed statistics for correlation are provided in **Supplementary Data 7 and 9**.

Defense islands are sometimes associated with MGEs integrated into distinct hotspots—regions of high genetic turnover—that may serve as catalysts for novel defensive strategies^9, 23^. The clustering of defense systems in these islands, especially within integrated MGEs, enhances the probability of horizontal co-transfer. If the repertoire of defense systems is primarily shaped by HGT, rather than functional synergies, we would expect the systems co-localizing in defense islands to correspond with those co-occurring in microbial genomes^23^. To explore the potential link between the co-occurrence and co-localization of defense systems, we selected 34 defense systems clustered in defense islands and analyzed the correlations among them.

Among the clustered systems, 105 pairs were identified with significant correlations (*p* < 0.05), which is lower than the 182 significant pairs observed when considering all defense systems. Notably, only one pair among the clustered systems exhibited a negative correlation **(Fig. 4a and Supplementary Data 9)**. Furthermore, there appears to be no obvious relationship between the number of defense systems clustered in defense islands and the number of significantly correlated pairs among those systems **(Fig. 4a and Supplementary Fig. 8)**. For instance, although RM and CRISPR-Cas systems were frequently found in defense islands, they did not exhibit an obvious higher number of correlations with other systems. In contrast, systems like BREX (13 pairs), Septu (13 pairs), and PT_PbeABCD (12 pairs), which were less commonly associated with defense islands, showed more significant correlations. These findings suggest that co-localization represents only a specific instance of co-occurrence, and the repertoire of defense genes is influenced by various factors beyond HGT.

The genomic context of defense islands supports the observed correlations among defense systems. For instance, the CBASS system frequently clusters within these islands, often flanked by defense genes related to RM and Abi systems **(Fig. 4b and Supplementary Data 10)**. These defense islands are not limited to defense genes alone; they also encompass a rich array of functional genes involved in transport, replication, recombination, repair, and transcription. This high genetic turnover within defense islands creates hotspots of genetic diversity, promoting the evolution of novel defense strategies in cold seep microorganisms^9^. Thus, the clustering and dynamic interactions within defense islands highlight their role in the co-localization and co-evolution of defense systems.

### Cold seep defense systems are abundant, actively expressed and site-dependent

To assess the distribution and activity of defense systems in cold seep prokaryotes, we calculated the relative gene abundance and transcript abundance of various defense genes and systems. In alignment with the prevalence of defense systems in microbial genomes **(Fig. 1a)**, RM, CRISPR-Cas, AbiE and SoFic systems exhibit higher metagenomic abundances, primarily due to the elevated abundances of restriction endonucleases (REases), methyltransferases (MTases), AbiEi, AbiEii, SoFic proteins, and Cas2 genes associated with these systems **(Fig. 5a and Supplementary Fig. 9)**. These differences are statistically significant at both systems and gene levels (*p* < 0.001). Although there were no significant differences in metatranscriptomic abundances among defense systems, the detection of their expression levels (averaging 3.64 transcripts per million [TPM], with over 1 TPM for 30 systems) indicate microbial defensive responses are active *in situ*. Additionally, most defense systems were detected across geographically diverse cold seep sediments (17 sites) at almost every sediment depth (0-68.55 meters below seafloor [mbsf]; **Supplementary Fig. 10)**, indicating a widespread presence of these systems in cold seeps globally. High abundances of defense systems were particularly notable in archaeal *Halobacteriota*, and bacterial *Desulfobacterota*, *Atribacterota*, and *Pseudomonadota* **(Fig. 5b)**. The relative abundance of defense systems varied considerably across phyla, with RM, CRISPR-Cas, AbiE and SoFic systems being the most predominant.

**Figure 5.**
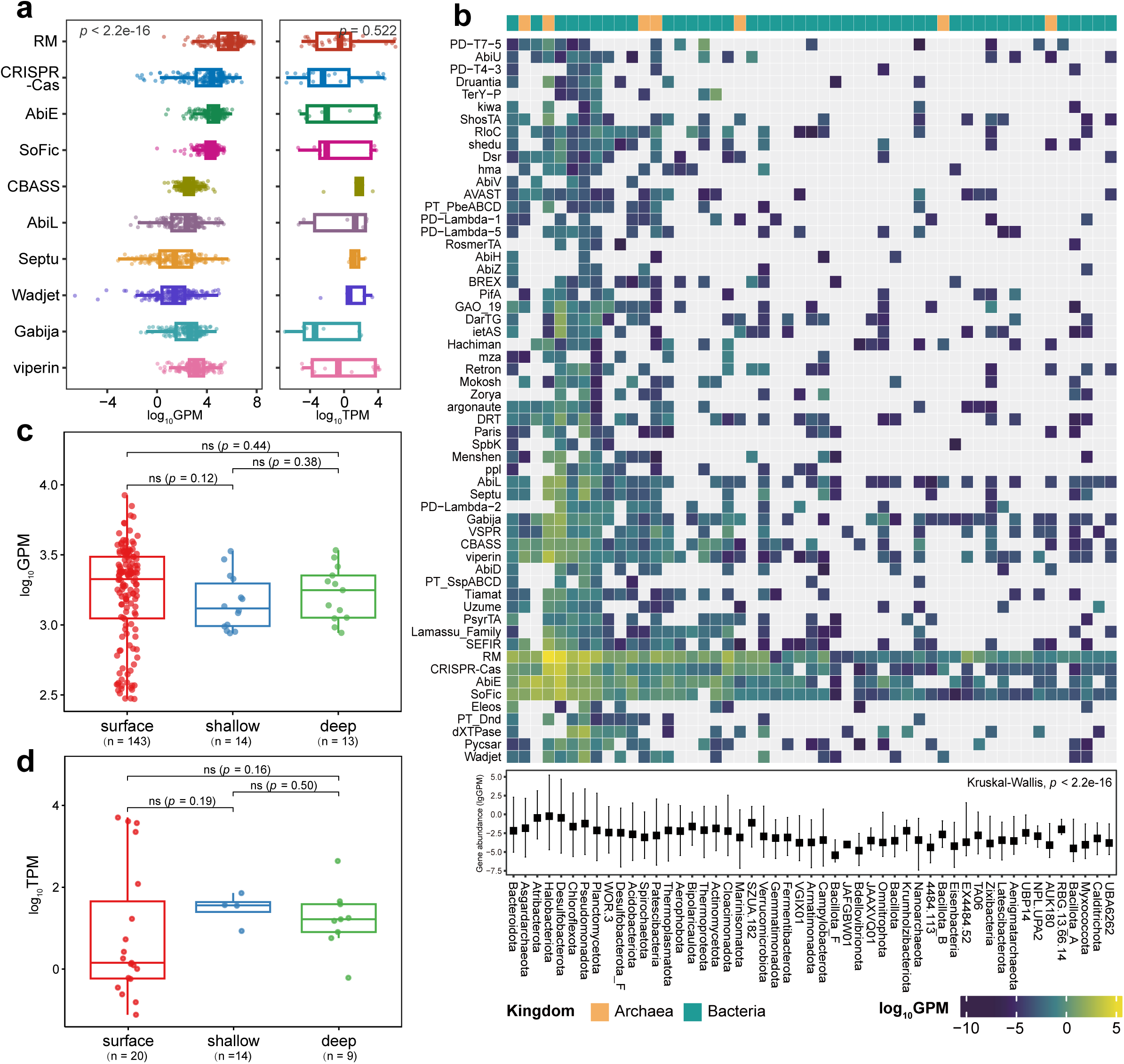
Relative abundance and expression of microbial defense systems in cold seep sediments. **(a)** Relative abundance (genes per million, GPM, n = 171, *p* < 2.2e-16) and transcript abundance (transcripts per million, TPM, n = 33, *p* = 0.522) of the top ten defense systems across different cold seep sediment samples. Different colors represent different types of defense systems. Statistical differences among defense system types were assessed using a two-sided Kruskal-Wallis rank-sum test. Boxplot: center line, median; box limits, upper and lower quartiles; whiskers, 1.5 × interquartile range; points, individual data values. **(b)** The heatmap displays the average gene abundance of various defense systems within each phylum in cold seep sediments, expressed in GPM. Only phyla with more than 20 defense systems are shown (n = 51). Gray squares indicate a GPM value of zero. Green labels represent bacterial phyla, while orange denotes archaeal phyla. The bar graph illustrates the average relative abundance of defense systems across phyla (*p* < 2.2e-16). Data are presented as mean ± SD. **(c)** Gene abundance of defense system genes at different sediment depths in cold seeps, measured in GPM. Sediment depths are categorized into three groups: surface (<1 meter below seafloor [mbsf], n = 143), shallow (1-10 mbsf, n = 14), and deep (>10 mbsf, n = 13). Differences were computed using a two-sided Wilcoxon rank-sum test. Boxplot: center line, median; box limits, upper and lower quartiles; whiskers, 1.5 × interquartile range; points, individual data values. **(d)** Expression levels of defense system genes at different sediment depths in cold seeps, measured in TPM. Sediment depths are categorized into three groups: surface (<1 mbsf, n = 20), shallow (1-10 mbsf, n = 4), and deep (>10 mbsf, n = 9). Differences were computed using a two-sided Wilcoxon rank-sum test. Boxplot: center line, median; box limits, upper and lower quartiles; whiskers, 1.5 × interquartile range; points, individual data values. Source data are provided as a Source Data file.

The distribution patterns of defense systems in global cold seep prokaryotes are shaped not only by host microbial taxa but also by their targets, such as viruses and MGEs. The distribution of cold seep microbes and viruses is influenced by sediment depth and site location^10, 44, 45^ ; consequently, these factors may also impact the distribution of defense systems. To assess potential depth stratification of defense systems, we categorized each metagenome based on depth ranges: surface (<1 mbsf), shallow (1 to 10 mbsf), and deep (>10 mbsf). Our analyses revealed no significant differences in the abundance of defense systems across these depth categories at either the metagenomic or metatranscriptomic level **(Fig. 5c, 5d and Supplementary Fig. 11)**. Significant differences in defense system abundance were found among different cold seep sites, with more pronounced differences in shallower depths (surface and shallow, *p* ≤ 0.05) compared to deeper depths (deep, *p* > 0.05; **Supplementary Fig. 12a**). Metatranscriptomic abundance also varied significantly across different sites, with a more notable difference observed in surface depths (*p* < 0.01; **Supplementary Fig. 12b**). Overall, these results suggest that the distribution of defense systems in cold seep prokaryotes is more strongly influenced by site-specific environmental and ecological factors than by sediment depth.

### Cold seep viruses employ diverse anti-defense genes to counteract host defense systems

We retrieved 13,336 single-contig viral genomes with estimated completeness of greater than 50% from 191 cold seep metagenomes **(Supplementary Fig. 13 and Supplementary Data 11)** using three virus identification tools: VIBRANT^46^, Virsorter2^47^ and geNomad^48^. These viral genomes were clustered into 9,971 species-level viral operational taxonomic units (vOTUs)^49^. Of these, 9,638 vOTUs (∼97%) could be taxonomically assigned, predominantly belonging to the *Duplodnaviria* phylum (n = 9,600), with a majority classified under the *Caudoviricetes* class (n = 9,599; **Supplementary Fig. 14 and Supplementary Data 12)**. However, less than 0.4% vOTUs (n = 36) within this class could be annotated further, highlighting an obvious knowledge gap in the taxonomy of deep-sea cold seep viruses^6, 10^.

We used the iPHoP v1.3.2 pipeline^50^ and CRISPR spacer sequence alignment to infer host-virus linkages, and identified a total of 2,688 connections between prokaryotes (810 genomes) and viruses (1,310 vOTUs; **Fig. 6a, Supplementary Fig. 15 and Supplementary Data 13)**. These host-virus linkages spanned 57 phyla, including 10 archaeal and 47 bacterial phyla. The most common predicted host phyla were *Chloroflexota* (n = 156), followed by *Halobacteriota* (n = 86), *Pseudomonadota* (n = 85), *Planctomycetota* (n = 56), and *Asgardarchaeota* (n = 52). Most of the identified connections involved viruses in the *Caudoviricetes* class infecting the *Chloroflexota* (731 pairs), and archaeal *Asgardarchaeota* (446 pairs) and *Halobacteriota* (284 pairs). The host-virus network revealed that the majority of clusters (60%, n=1,616) consisted of more than one viral or prokaryotic genome. This suggests that a single virus could infect multiple prokaryotic populations, or a single prokaryote may be susceptible to several viral population^51^, indicating a complex relationship between viruses and prokaryotes in cold seeps.

**Figure 6.**
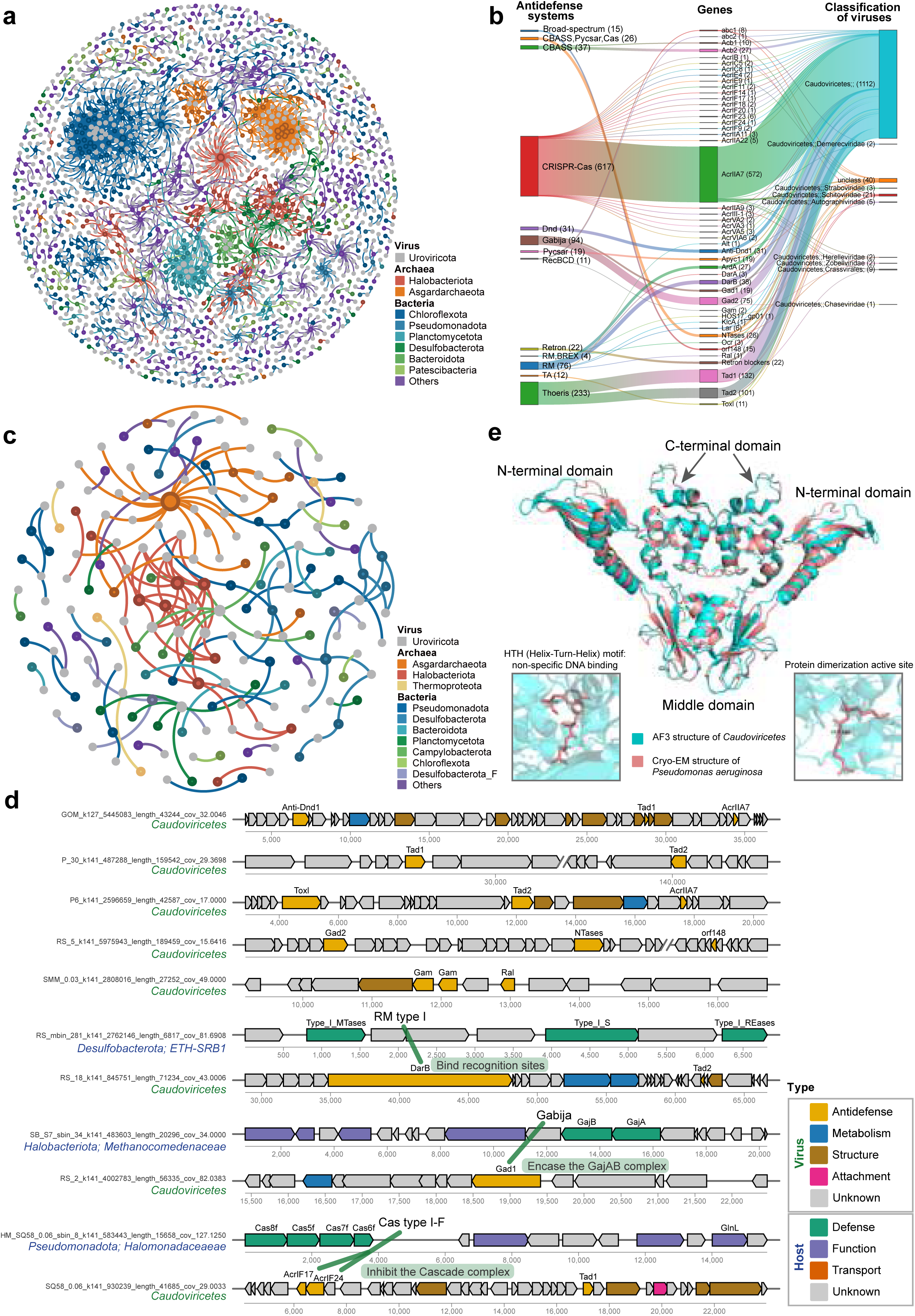
Interactions between cold seep viral anti-defense systems and host defense systems. **(a)** The network diagram illustrates 2,688 host-virus pairs identified in cold seeps. Prokaryotes are colored according to their phyla. **(b)** Anti-defense genes identified in cold seep viruses, showing the types of anti-defense systems and corresponding viral taxonomy. **(c)** Illustration of 155 host-virus pairs where host prokaryotes contain defense genes and the corresponding viruses possess anti-defense genes. **(d)** Genomic organization of viral gene clusters containing multiple types of anti-defense systems, along with the corresponding host defense systems. Viral genes are categorized and color-coded: anti-defense genes (yellow), metabolic genes (blue), structural genes (brown), attachment genes (pink), and unknown genes (grey). Host defense genes are highlighted in green, transport-related genes in orange, and other functional genes in purple. Green lines indicate interactions between host defense systems and viral anti-defense systems. **(e)** Structural comparison of the AcrIF24 protein dimer from a *Caudoviricetes* virus infecting *Halomonas* sp. (blue) with the known cryo-electron microscopy structure from *Pseudomonas aeruginosa* (pink, PDB code: 7ELM). A detailed view highlights key sites involved in AcrIF24 protein dimerization and binding to the Csy complex.

Viruses encode an arsenal of anti-defense proteins that facilitate infection by disabling various prokaryotic defense mechanisms^18, 24^. A total of 1,197 anti-defense genes were detected from 11% viral genomes (n = 1,065), predominantly comprising anti-CRISPR-Cas, anti-Thoeris, anti-Gabija, and anti-RM genes (**Fig. 6b and Supplementary Data 14)**. While most anti-defense genes were detected across a broad range of viruses, some were found only in specific viral genomes. For example, anti-CRISPR, anti-RM, anti-Thoeris, and anti-Gabija genes were commonly found, whereas anti-RecBCD genes and broad-spectrum anti-defense genes (*orf148* ^52^, a broad-spectrum anti-defense gene in *Escherichia* phage OSYSP that targets conserved host barrier defenses and metabolic pathways to weaken the host’s multi-layered antiviral mechanisms) were exclusively detected in certain *Caudoviricetes* viruses. From the 2,688 host-virus linkages, we identified 155 pairs in which host microbes contained defense genes while the corresponding viruses possessed anti-defense genes **(Fig. 6c and Supplementary Data 13)**. These connections were predominantly observed between viruses in the *Caudoviricetes* class and their hosts, including archaeal groups such as *Asgardarchaeota* (38 pairs) and *Halobacteriota* (26 pairs), as well as various bacterial taxa including *Pseudomonadota* (25 pairs) and *Desulfobacterota* (24 pairs). Notably, a single virus employing two anti-defense systems could infect two ETH-SRB1 genomes belonging to the *Desulfobacterota* phylum, and one *Asgardarchaeota* genome was found to harbor six defense systems to fend off infection by five viruses **(Fig. 6c and Supplementary Data 13)**. These findings suggest that the distribution of defense and anti-defense genes appear lineage dependent, particularly among various archaeal lineages and their viruses^10, 25, 53^.

A single virus may encode multiple anti-defense genes, enabling it to counter various host defense strategies or adapt to different defenses employed by the same host throughout its life cycle^25^. We identified 118 viruses that encode two or more types of anti-defense systems **(Supplementary Data 13)**. For instance, a viral genome from the *Caudoviricetes* family contains AcrIF17, AcrIF24, and Tad1 genes to counteract the CRISPR-Cas and Thoeris defense systems of its host belonging to the phylum *Pseudomonadota* **(Fig. 6c)**. Additionally, two viral genomes from the same family harbor DarB and Tad2 genes to counteract the RM and Thoeris defense systems of their host, ETH-SRB1 **(Fig. 6d and Supplementary Data 13 in green)**. The predicted dimeric structure of AcrIF24 from *Caudoviricetes* exhibits high similarity (TM-score = 0.86) to the anti-CRISPR protein form *Pseudomonas aeruginosa* **(Fig. 6e)**, which has been experimentally confirmed to inhibit Cas7f^54^. AcrIF24 interacts with immune system components through steric or allosteric effects in a stoichiometric, non-enzymatic manner^54^. Many other anti-defense genes that function similarly were also detected in cold seeps, including most Acrs (e.g., AcrIIA7, AcrIIA2, AcrIIA4, AcrIIIB1, and AcrIIC1). In contrast, enzymatic inhibitors, which effectively suppress host defenses by permanently neutralizing microbial immunity^24, 55, 56^, were also identified, such as Acb1, NTases, AcrIII-1, and AcrVA1-5. Additionally, several anti-defense systems with unknown molecular mechanisms, such as Anti-Dnd, Gad2, Abc1, and KlcA, were also present. Overall, viral anti-defense systems engage in complex evolutionary interactions with host defense systems, employing enzymatic, non-enzymatic, and unknown molecular mechanisms in cold seep ecosystems.

### Defense systems protect key metabolic microbial groups from viral infections

Anaerobic methane oxidation, sulfate reduction, and nitrogen fixation, associated with the activities of the *mcrA*, *dsrA*, and *nifH* genes, respectively, are key metabolic processes in cold seeps^57–59^. The significant correlation between the abundance of defense systems and these key metabolic genes **(Supplementary Fig. 16)** underscores the role of defense systems in protecting functional microbes from viral and MGE infections. To further explore this relationship, we analyzed 35 anaerobic methanotrophic archaea (ANME), 47 sulfate-reducing bacteria (SRB), and 57 diazotrophs identified from 3,813 species-level MAGs. The total number of systems and the diversity of families varied significantly across individual microbial groups, particularly among SRB and diazotrophs **(Fig. 7b)**. While the overall repertoire of defense systems remain relatively conserved across major microbial groups **(Fig. 7a, Supplementary Fig. 17, and Supplementary Data 15)**, their abundance and diversity differ markedly among groups, underscoring the complexity of defense system distribution across different microbial groups. For example, ANME generally encode fewer defense systems, with a limited diversity of types. Within SRB, ETH-SRB1 encode on average 17.6 defense systems across 7.3 types, whereas another SRB strain in *Gemmatimonadota* order (SY6_cobin_101) encodes only 3 systems of just 1 type (belonging to AbiE; **Fig. 7b and Supplementary Data 15**).

**Figure 7.**
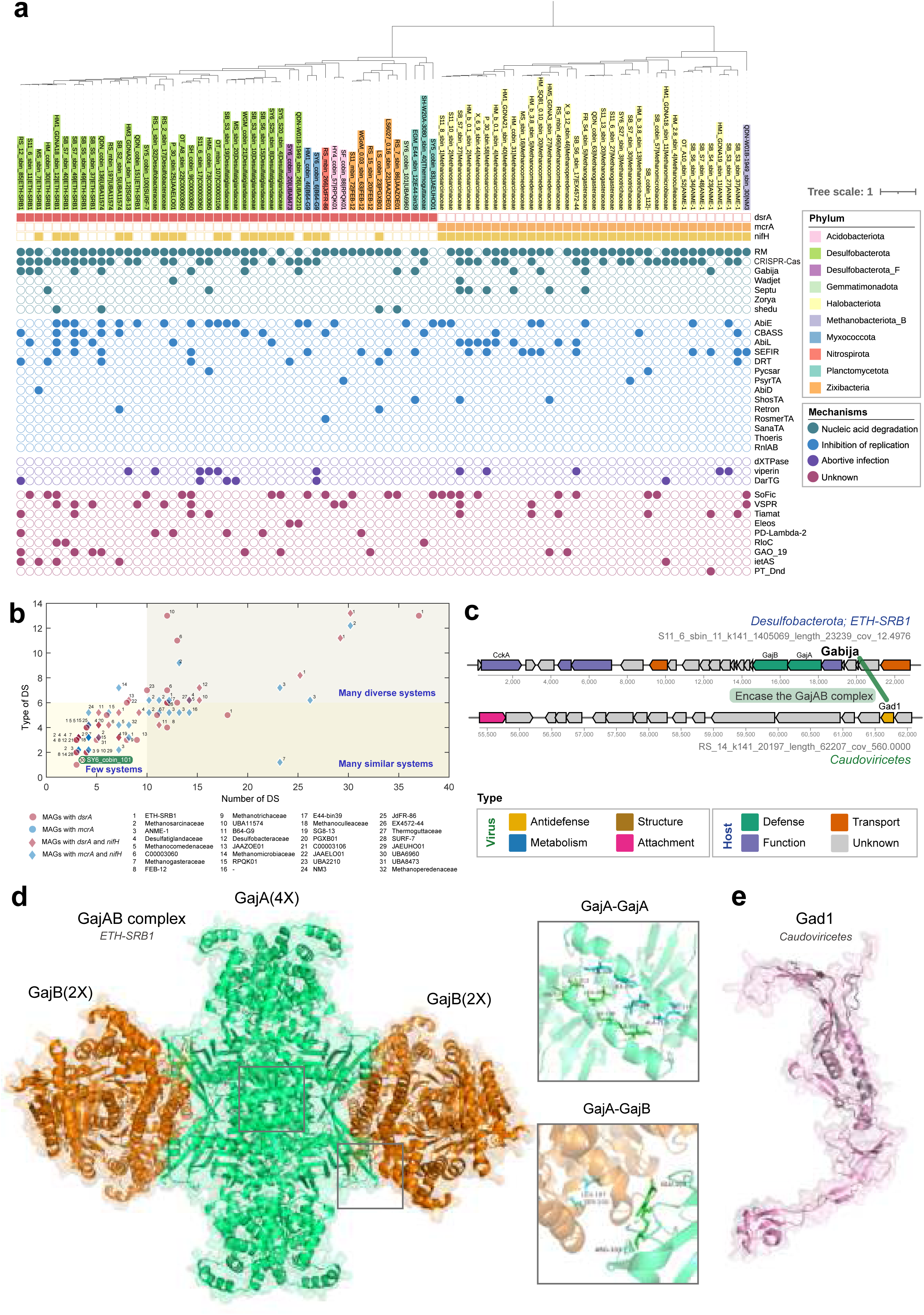
Defense competition between key metabolic microbial groups in cold seeps and their viruses. **(a)** Type of defense systems in key metabolic microbial groups, including anaerobic methanotrophic archaea (ANME), sulfate-reducing bacteria (SRB), and diazotrophs. Colored squares indicate the presence of key functional genes: *mcrA* for ANME, *dsrA* for SRB, and *nifH* for diazotrophs. Colored circles represent the types of defense systems detected in these microbial groups. Different phyla are denoted by distinct colors, and different defense mechanisms are indicated by circles of various colors. To address potential overlap, regions where *mcrA* and *nifH* genes coexist have been highlighted, reflecting functional versatility within the same microenvironments. **(b)** Relationship between the number of defense systems and the diversity of defense types in key metabolic microbial groups, categorized into three groups based on quantity: few systems, diverse systems, and similar systems. **(c)** Genomic organization of the Gabija defense system and the viral *gad1* gene in a virus-host pair involving ETH-SRB1 and a *Caudoviricetes* virus. Genes are color-coded: anti-defense genes (green), structural genes (purple), metabolic genes (pink), attachment genes (orange), transport genes (yellow), defense genes (blue), and genes of unknown function (gray). Green lines indicate interactions between the host defense system and the viral anti-defense system. **(d)** The structure of GajAB complex in cold seep ETH-SRB1 predicted by Alphafold3, including detailed diagrams of the main interaction sites between GajA proteins and between GajB proteins. **(e)** The structure of Gad1 protein in a cold seep *Caudoviricetes* virus predicted by Alphafold3.

In detail, SRB predominantly encode type IIG and type II RM systems, followed by type I RM systems. ANME and diazotrophs primarily encode type IIG and type I systems, with type II RM systems following (**Supplementary Fig. 18 and Supplementary Data 15)**. The number of these systems in ANME is obvious smaller compared to the other two functional groups. RM systems primarily function in the early stages of viral infection to eliminate invading nucleic acids^11^, indicating that cold seep prokaryotes possess the ability to respond rapidly and clear viruses promptly. CRISPR-Cas systems are categorized into two classes and six types based on the set of Cas genes they encode and whether their effector is a single protein or a protein complex^13, 60, 61^. Both classes of CRISPR-Cas were present in cold seep microbes, with class I being more abundant in both number and diversity (**Supplementary Fig. 19 and Supplementary Data 15)**. Most archaea, including ANME, primarily encode type I CRISPR-Cas systems, while the majority of bacteria, including SRB and diazotrophs, mainly encode both type I and type III CRISPR-Cas systems. CRISPR-Cas has been shown to contribute to shaping a highly dynamic network of interactions between viruses/MGEs and their prokaryotic hosts in various ecosystems^62–64^.

Additionally, SRB tend to harbor two or more Gabija systems, particularly in ETH-SRB1 strains, with some encoding as many as four Gabija systems (**Supplementary Fig. 20 and Supplementary Data 15)**. Overall, these findings highlight the diversity and complexity of defense systems across various microbial groups, likely shaped by distinct evolutionary pressures and ecological niches^19^.

Gabija is effective in providing bacterial immunity against certain bacteriophages, as shown by its strong defense against phi29, rho14, phi105, and SpBeta in *Bacillus cereus* VD045 ^7^. Through sequence alignment and both sequence- and structure-based phylogenetic analyses **(Supplementary Fig. 21)**, we found that Gabija is widely present in cold seep microbes, spanning 45 phyla (including 9 archaeal and 36 bacterial phyla) and encompassing the three key metabolic microbial groups. We discovered that one ETH-SRB1 strain encodes two Gabija systems, and the virus infecting it carries the *gad1* anti-defense gene **(Fig. 7c)**. This ETH-SRB1 strain harbored both a complete and an incomplete Gabija system. In the complete system, the GajA protein contains an N-terminal ATPase domain and a C-terminal Toprim (topoisomerase-primase) domain, while the GajB protein includes an AAA domain and a UvrD-like helicase^30, 65^. In contrast, the GajB of the incomplete Gabija system lacks the UvrD-like helicase **(Supplementary Fig. 20)**. The predicted structure shows similarity (TM-score = 0.70) to the Gabija protein from *Bacillus cereus* VD045^30^ **(Fig. 7d)**, suggesting potential conservation of functional domains. The viral protein Gad1 **(Fig. 7e)**, larger than other phage immune evasion proteins (∼35 kDa), appears to bind and encapsulate the GajAB complex, potentially blocking the degradation of target DNA. The binding of Gad1 to the GajAB complex encoded by a virus targeting ETH-SRB1 suggests that Gad1 may inhibit the Gabija system in this strain.

Considering the differences in the number and types of defense systems among functional microbial groups, we hypothesize that the specific pairs of co-occurring and negatively associated defense systems vary across these groups. As anticipated, defense systems that co-occurred in one group were absent in another **(Supplementary Fig. 22 and Supplementary Data 16)**. For example, AbiE system in SRB exhibited 28 co-occurrences, including with CBASS, DarTG, Gabija, and Mokosh, none of which were observed in ANME or diazotrophs. These findings highlight the unique defensive strategies employed by different microbial lineages. Previous research has revealed that the relationships between defense systems likely depend on environmental and genetic factors that select for a particular anti-virus immunity strategy, and that synergistic immunity provides an evolutionary advantage to microbial populations^23^. Overall, our results indicate that cold seep microbial defense systems are generally mechanistically compatible, allowing microbes to adopt diverse and flexible strategies for anti-virus defense based on their unique environmental and genetic contexts.

### Implications of microbe-virus relationships in cold seeps

Cold seep ecosystems, characterized by the seepage of hydrocarbon-rich fluids into deep-sea sediments, harbor a diverse array of previously uncharacterized microbes and viruses engaged in a continuous evolutionary arms race. In this study, we identified diverse defense systems in cold seep microbes, including a substantial proportion of candidate systems. These defense systems frequently co-occur and may work synergistically to combat infections from MGEs and viruses. Experimental validation confirmed that several of these candidate defense systems effectively protect against viral infections. Notably, defense systems such as RM, CRISPR-Cas, AbiE, and SoFic exhibited high metagenomic abundances, widespread distribution, and active expression across various microbial genomes from different sediment depths and geographical locations, underscoring their vital role in microbial survival within these extreme environments. In response, cold seep viruses have evolved diverse anti-defense mechanisms, functioning through enzymatic, non-enzymatic, and as-yet-unknown mechanisms to evade host defenses. The presence of multiple viral anti-defense genes within single viral genomes, along with various defense genes in their hosts, highlights the intricate and dynamic interactions between these viruses and their microbial hosts, reflecting a potential ongoing evolutionary arms race. Functionally critical lineages in cold seeps, such as ANME, SRB, and diazotrophs, adjust the correlations among different defense systems according to their ecological niches and environmental pressures. Overall, these findings emphasize the complexity and flexibility of interactions within microbial defense systems in extreme environments. The ability of cold seep microbes to maintain diverse and adaptable defense mechanisms highlights their resilience and evolutionary success in one of Earth’s most challenging habitats. This study deepens our understanding of virus-microbe interactions in cold seep ecosystems and underscores the significance of microbial defense systems in shaping microbial ecology and evolution in extreme environments.

## Methods

### Dataset and processing

Metagenomic assemblies were generated from 171 deep-sea sediment samples collected from 17 geographically diverse cold seep sites, with sediment depths ranging from 0 to 68.55 mbsf **(Supplementary Fig. 1)**. The construction of the genomic inventory followed the methods detailed in our previous study^5^. Raw metagenomic sequence data underwent quality control and were assembled into contigs sequences. Contigs exceeding 1,000 base pairs in length were selected for binning. The resulting MAGs were deduplicated using dRep v3.4.0 ^66^ (parameters: - comp 50 -con 10) at a 95% average nucleotide identity threshold, yielding a total of 3,813 cold seep prokaryotic MAGs. Initial taxonomic assignment of each MAG was conducted using GTDB-TK v2.1.3 ^67^ with reference to the Genome Taxonomy Database (GTDB) release R214, followed by validation through a maximum-likelihood phylogenomic tree. For the archaeal MAGs, the phylogenomic tree was inferred based on the concatenation of 53 conserved single-copy genes as marker genes, while for the bacterial MAGs, it was based on 120 marker genes. The trees were constructed using IQ-TREE v2.2.0.3 (parameters: -m MFP -B 1000)^68^ and visualized by using iTOL v6 ^69^. Sankey plots were constructed using Pavian (https://fbreitwieser.shinyapps.io/pavian/)^70^. Protein-coding sequences were predicted from the MAGs using Prodigal v2.6.3 (parameter: -p meta)^71^, resulting in a gene catalog annotated from all MAGs.

Thirty-three metatranscriptomes, originating from cold seeps of the South China Sea in regions of Jiaolong, Haima, Qiongdongnan Basin, and the Shenhu area^59, 72–74^ **(Supplementary Fig. 1)**, were subjected to quality control using the Read_QC module of the metaWRAP v1.3.2 pipeline (parameters: --skip-bmtagger)^75^. Following this, SortMeRNA v2.1 was utilized to remove ribosomal RNA^76^. The resulting clean reads were then aligned to the gene catalog annotated from all MAGs, employing Salmon v.1.9.0 (parameters: -validateMappings -meta)^77^, to quantify the transcript abundance of prokaryotic defense genes. Transcript levels were normalized to transcripts per million (TPM).

### Identification of defense genes, systems, and islands

Defense genes and systems were identified using DefenseFinder v1.2.1 (parameters: --db-type gembase)^19^ and PADLOC v2.0.0 (parameter: --fix-prodigal)^33^. DefenseFinder detects 710 genes related to 154 defense system families by matching sequences to 845 Hidden Markov Models (HMMs) representing experimentally validated antiviral proteins. It classifies defense systems based on predefined genetic architecture rules, which require mandatory proteins to be genomically adjacent and exclude prohibited components. PADLOC detects 385 defense subsystems using HMM libraries and classification files, enforcing gene proximity via the max_separation parameter and validating associations through co-occurrence frequency statistics (one-sample proportion test, *p* < 0.001). Candidate defense systems were annotated according to the PADLOC database, which includes HECs (Hma-Embedded Candidates) and PDCs (Phage Defense Candidates)^33^. These candidates are predicted using a “guilt-by-embedding” strategy, where uncharacterized genes are recurrently embedded within known defense systems (e.g., Hma, RM, BREX, DISARM)^34^. The classification of proteins within a single defense system was determined based on parameters established by DefenseFinder and PADLOC, including maximum intergenic distances, domain architecture consistency, and functional annotation rules. Results from the two tools were merged, with duplicate results compared based on e-values to retain those with higher confidence. Defense islands were defined as arrays of defense genes (or defense systems) separated by no more than ten genes and containing at least five genes from at least three different defense systems^9^.

Gene abundances of gene catalog annotated from all MAGs were quantified using Salmon v.1.9.0 in mapping-based mode (parameters: --meta --validateMappings)^77^ and read counts were normalized to GPM (genes per million). The abundance of a defense system is represented by the sum of the abundances of the genes within that system divided by the average number of genes in the system.

### Identification of viruses and virus-host linkages

Potential viral sequences were identified from metagenomic assemblies (contig lengths greater than 10 kb) derived from the aforementioned 171 cold seep site samples and an additional 20 cold seep samples. Three methodologies were used: geNomad v1.7.0 (parameters: end-to-end)^48^, Virsorter2 v2.2.3 (parameters: default)^47^, and VIBRANT v1.2.1 (parameters: default)^46^. The completeness and contamination of these sequences were assessed using CheckV v1.0.1 with the database v1.5 ^78^. Sequences with an estimated completeness of 50% or more were selected for clustering into species-level viral operational taxonomic units (vOTUs). Clustering parameters followed CheckV guidelines: 95% average nucleotide identity and a minimum aligned fraction of 85%. This process resulted in the identification of 9,971 species-level vOTUs. Taxonomic assignment of these vOTUs was conducted using geNomad v1.7.0. The resulting taxonomic lineages adhere to the classification outlined in the International Committee on Taxonomy of Viruses (ICTV) Virus Metadata Resource (VMR) number 19. Open reading frames (ORFs) within each vOTU were predicted using Prodigal v2.6.3 (parameter: -p meta)^71^. Functional annotation of the viral sequences was performed utilizing the virome model in DRAM v1.4.0 ^79^.

Linkages between viruses and their hosts were primarily determined using the iPHoP v1.3.2 pipeline^50^, utilizing a database constructed from 3,813 prokaryotic MAGs identified in this study and the iPHoP_db_Sept21_rw. iPHoP employs a variety of methods to predict virus-host linkages, including Blastn v2.12.0 ^80^ alignment of viral genomes against the iPHoP_db_Sept21_rw database and its CRISPR spacer database; The k-mer composition similarity between viral and host genomes was analyzed using four methods: RaFAH v0.3 ^81^, WIsH v1.0 ^82^, VirHostMatcher-Net^83^, and PHP^84^. In addition to the iPHop methods, for contigs identified with CRISPR-Cas systems, CRISPRCasFinder v4.2.21 ^85^ was used to identify CRISPR arrays flanking the Cas genes. Local alignments of extracted spacers (with lengths greater than 25 base pairs) against viral genomes were performed using Blastn. Only BLAST matches with 100% alignment coverage and at most two mismatches were considered high-confidence matches. The network graph between hosts and viruses was visualized using Gephi v0.9.2 ^86^. The workflow for this part was illustrated using Draw.io v23.1.5 (https://app.diagrams.net/).

### Identification of anti-defense genes

The anti-defense genes of viruses were primarily identified through two approaches. (1) APIS database search. We utilized the Anti-Prokaryotic Immune System (APIS) database^87^, which contains experimentally validated viral proteins that counteract prokaryotic immune systems and their associated protein families. All viral proteins from our dataset were compared against the APIS database using DIAMOND blastp v2.0.8 (parameters: -e 1e-10 --id 30)^88^ and HMMER v3.3.2 (parameters: -E 1e-10). (2) Reference sequence comparison. We compiled sequences of viral anti-defense proteins as documented by Mayo-Muñoz et al. in their review^24^. These 36 reference sequences were downloaded from the NCBI database and used to construct a custom database using the DIAMOND makedb command. Our viral proteins were then compared against this database using DIAMOND BLASTp v2.0.8 (parameters: -e 1e- 10 --id 30). The results were obtained using two methods, with the more reliable results selected based on e-values to derive the final set of anti-defense genes.

### Experimental validation of candidate defense systems

Coding sequences of the candidate defense systems were commercially synthesized and cloned into the pQE82L expression vector. The recombinant vectors were transformed into *E. coli* B (ATCC^®^ 11303™). Cells transformed with the empty vector served as controls. Plaque assays were conducted following established protocols^14, 89^. A single colony from a fresh LB agar plate was inoculated into LB broth containing ampicillin (100 µg/mL) and grown at 37°C to an OD_600_ of approximately 0.6. Protein expression was induced by adding 0.2 mM isopropyl β-D-1-thiogalactopyranoside (IPTG). After an additional hour of incubation, 500 µL of the bacterial culture was mixed with 14.5 mL of 0.5% LB top agar, and the mixture was poured onto LB plates containing ampicillin (100 µg/mL) and IPTG (0.1 mM). The plates were then spotted with 4 µL aliquots of phage dilutions prepared in LB medium. Eight 10-fold serial dilutions were used for each phage: dilutions 10^-1^ to 10^-8^ for phage T7, and 10^0^ to 10^-7^ for phages T4 and T5. Plates were incubated overnight at 25°C before imaging. All the assays are repeated at least 3 times with similar results.

### Functional annotations

Functional annotation of MAGs was performed by searching against the KEGG, Pfam, MEROPS, and dbCAN databases using DRAM v1.4.0 (parameters: --min_contig_size 1000)^79^. Gene context was visualized with Chiplot (https://www.chiplot.online/).

Plasmid and provirus sequences were identified from prokaryotic MAGs using geNomad v1.7.0 ^48^. To classify different types of MGEs, such as IS elements and transposons (IS_Tn), integrons, and conjugative elements (CE), recombinase marker genes, were identified through alignment with the proMGE database^90^ using HMMER v3.3.2. For the identification of key metabolic genes in cold seep prokaryotes, *dsrA* genes were detected using DiSco v1.0.0 ^91^, and *mcrA* genes were identified using METABOLIC v4.0 ^92^ and DRAM v1.4.0 ^79^. Nitrogen fixation marker genes *nifH* genes were detected using three methods: (1) protein sequences in the MAGs were compared against NCycDB^93^ using DIAMOND blastp v2.0.8 ^88^, (2) potential *nifH* sequences were searched based on *nifH* reference sequences (n = 1,271) from the Greening lab metabolic marker gene database using DIAMOND blastp v2.0.8 (parameters: --id 65)^88^, and (3) *nifH* genes were extracted using HMMER v3.3.2 with the TIGRFAM model TIGR01287.1.

### Protein sequence and structural analysis

To verify the accuracy and phylogenetic placement of the identified defense and anti-defense genes, multiple sequence alignments were performed using MAFFT v7.505 ^94^. Alignments were trimmed using TrimAl v1.4.1 under default settings^95^. Maximum likelihood phylogenetic trees were constructed using IQ-TREE v2.2.0.3 (parameters: - m MFP -B 1000)^68^. Trees were visualized and annotated using iTOL v6 ^69^. Protein domains were identified using the Batch Protein Annotation plugin in Tbtools-II v2.042 ^96^, which integrates results from InterProScan^97^. Sequence conservation among proteins was aligned and visualized using Jalview v2.11.3.3 ^98^. Structural predictions for proteins, including GajA and GajB, were performed using ColabFold v1.5.5 which leverages AlphaFold2^99–101^ on September 14, 2023. Structures of the AcrIF24 dimer, GajAB octamer, and Gad1 protein were predicted using AlphaFold3^102^ on August 30, 2024. The predicted structures were evaluated based on pLDDT and Predicted Aligned Error (PAE) scores **(Supplementary Fig. 23)**. The reference structures AcrIF24 dimer and GajAB octamer were obtained from experimentally determined Csy-AcrIF24 complex (PDB code: 7ELM) and GajAB-Gad1 complex (PDB code: 8U7I). Homologous and structurally similar proteins were identified using the Foldseek web interface^103^. Protein structures were visualized and aligned using PyMOL v2.5.5 ^104^.

### Statistical analyses

Statistical analyses were conducted using R v4.2.3. Normality of the data was assessed using the Kolmogorov-Smirnov test, and homogeneity of variances was tested using Levene’s test. The Wilcoxon rank-sum test was employed to compare characteristics of defense systems between archaeal and bacterial groups. For comparisons involving more than two groups—such as defense system and gene densities among different microbial phyla, or differences in gene and transcript abundances among various types of defense systems or genes—the Kruskal-Wallis test was used. Pearson correlation analysis was conducted using R’s default method (cor.test function with Pearson method) to evaluate relationships between defense systems and their subsystems, based on abundance profiles. All correlation analyses were performed on log_10_-transformed abundance data to meet normality assumptions. For multiple comparisons, *p*-values were adjusted using the Benjamini-Hochberg false discovery rate (FDR) correction with a significance threshold of 0.05.

## Data availability

The non-redundant MAGs catalog has been deposited in the NCBI database under BioProject PRJNA950938. Sequences of defense systems, viral genomes, viral anti-defense systems, and phylogenetic trees of GajAB based on amino acid sequences and protein structures, generated in this study, are available at https://doi.org/10.6084/m9.figshare.27004438. All additional data supporting the findings of this study are provided within the article and its Supplementary Information Files. Source data are provided with this paper.

## Code availability

The present study did not generate codes, and mentioned tools used for the data analysis were applied with default parameters unless specified otherwise.

## Acknowledgements

The work was supported by National Key R&D Program of China (No. 2024YFC2816200 to X.D.), National Science Foundation of China (No. 92351304 to X.D., No. 42376115 to X.D., No. 42406109 to Y.H., No. 32470036 to R.C. and No. 32100025 to R.C.), Natural Science Foundation Project of Xiamen City (No. 3502Z202373076 to X.D.), Natural Science Foundation of Fujian Province (No. 2023J06042 to X.D.), Scientific Research Foundation of Third Institute of Oceanography, MNR (No. 2022025 to X.D., No. 2023022 to X.D. and No. 2025013 to Y.H.), Open Funding Project of State Key Laboratory of Microbial Metabolism (No. MMLKF23-05 to X.D.) and Fundamental Research Funds for the Central Universities (No. 2662025SYPY003 to R.C.).

## Author contributions

X.D. conceived and designed the study. Y.H. and J.L. performed the omics analyses. C.L. contributed to viral identification and classification. R.C. designed and conducted the experimental implementations. X.L. participated in the analysis of some metagenomic data. C.L., F.X., J.P. and W.X. actively participated in discussions and data interpretation. F.W. and H.J. provided valuable suggestions for manuscript revision. Y.H., J.L. and X.D. wrote the manuscript with input from all authors.

## References

1. Dong, X. et al. Metabolic potential of uncultured bacteria and archaea associated with petroleum seepage in deep-sea sediments. Nature Communications 10, 1816 (2019).

2. Joye, S.B. The geology and biogeochemistry of hydrocarbon seeps. Annual Review of Earth and Planetary Sciences 48, 205–231 (2020).

3. Dong, X. et al. Thermogenic hydrocarbon biodegradation by diverse depth-stratified microbial populations at a Scotian Basin cold seep. Nature Communications 11, 5825 (2020).

4. Kleindienst, S. et al. Diverse sulfate-reducing bacteria of the *Desulfosarcina/Desulfococcus* clade are the key alkane degraders at marine seeps. The ISME Journal 8, 2029–2044 (2014).

5. Yang, X.Y., Shen, Z., Lin, Q. & Fu, T.M. Molecular basis of Gabija anti-phage supramolecular assemblies. Nature 15, 836 (2023).

6. Li, Z. et al. Deep sea sediments associated with cold seeps are a subsurface reservoir of viral diversity. The ISME Journal 15, 2366–2378 (2021).

7. Doron, S. et al. Systematic discovery of antiphage defense systems in the microbial pangenome. Science 359, eaar4120 (2018).

8. Millman, A. et al. An expanded arsenal of immune systems that protect bacteria from phages. Cell Host & Microbe 30, 1556–1569 (2022).

9. Beavogui, A. et al. The defensome of complex bacterial communities. Nature Communications 15, 2146 (2024).

10. Peng, Y. et al. Viruses in deep-sea cold seep sediments harbor diverse survival mechanisms and remain genetically conserved within species. The ISME Journal 17, 1774–1784 (2023).

11. Georjon, H. & Bernheim, A. The highly diverse antiphage defence systems of bacteria. Nature Reviews Microbiology 21, 686–700 (2023).

12. Tock, M.R. & Dryden, D.T.F. The biology of restriction and anti-restriction. Current Opinion in Microbiology 8, 466–472 (2005).

13. Hille, F. et al. The biology of CRISPR-Cas: backward and forward. Cell 172, 1239–1259 (2018).

14. Cheng, R. et al. Prokaryotic Gabija complex senses and executes nucleotide depletion and DNA cleavage for antiviral defense. Cell Host & Microbe 31, 1331–1344 (2023).

15. Rousset, F. et al. A conserved family of immune effectors cleaves cellular ATP upon viral infection. Cell 186, 3619–3631 (2023).

16. Lopatina, A., Tal, N. & Sorek, R. Abortive infection: bacterial suicide as an antiviral immune strategy. Annual Review of Virology 7, 371–384 (2020).

17. Saenz, J.S., Seifert, J. & Rios-Galicia, B. Antiviral defense systems in the rumen microbiome. mSystems 10, e01521–24 (2025).

18. Piel, D. et al. Phage–host coevolution in natural populations. Nature Microbiology 7, 1075–1086 (2022).

19. Tesson, F. et al. Systematic and quantitative view of the antiviral arsenal of prokaryotes. Nature Communications 13, 2561 (2022).

20. Bernheim, A. & Sorek, R. The pan-immune system of bacteria: antiviral defence as a community resource. Nature Reviews Microbiology 18, 113–119 (2020).

21. Rocha, E.P.C. & Bikard, D. Microbial defenses against mobile genetic elements and viruses: Who defends whom from what? PLoS Biology 20, e3001514 (2022).

22. Picton, D.M. et al. The phage defence island of a multidrug resistant plasmid uses both BREX and type IV restriction for complementary protection from viruses. Nucleic Acids Research 49, 11257–11273 (2021).

23. Wu, Y. et al. Bacterial defense systems exhibit synergistic anti-phage activity. Cell Host & Microbe 32, 557–572.e556 (2024).

24. Mayo-Muñoz, D., Pinilla-Redondo, R., Camara-Wilpert, S., Birkholz, N. & Fineran, P.C. Inhibitors of bacterial immune systems: discovery, mechanisms and applications. Nature Reviews Genetics 25, 237–254 (2024).

25. Hampton, H.G., Watson, B.N.J. & Fineran, P.C. The arms race between bacteria and their phage foes. Nature 577, 327–336 (2020).

26. Duan, N., Hand, E., Pheko, M., Sharma, S. & Emiola, A. Structure-guided discovery of anti-CRISPR and anti-phage defense proteins. Nature Communications 15, 649 (2024).

27. Gussow, A.B. et al. Machine-learning approach expands the repertoire of anti-CRISPR protein families. Nature Communications 11, 3784 (2020).

28. Serfiotis-Mitsa, D. et al. The structure of the KlcA and ArdB proteins reveals a novel fold and antirestriction activity against Type I DNA restriction systems in vivo but not in vitro. Nucleic Acids Research 38, 1723–1737 (2010).

29. Hobbs, S.J. et al. Phage anti-CBASS and anti-Pycsar nucleases subvert bacterial immunity. Nature 605, 522–526 (2022).

30. Antine, S.P. et al. Structural basis of Gabija anti-phage defence and viral immune evasion. Nature 625, 360–365 (2024).

31. An, L. et al. Global diversity and ecological functions of viruses inhabiting oil reservoirs. Nature Communications 15, 6789 (2024).

32. Zhong, Z.-P. et al. Viral potential to modulate microbial methane metabolism varies by habitat. Nature Communications 15, 1857 (2024).

33. Payne, L.J. et al. Identification and classification of antiviral defence systems in bacteria and archaea with PADLOC reveals new system types. Nucleic Acids Research 49, 10868–10878 (2021).

34. 34. Payne, L.J., Hughes, T.C.D., Fineran, P.C. & Jackson, S.A. New antiviral defences are genetically embedded within prokaryotic immune systems. bioRxiv, 10.1101/2024.01.29.577857 (2024).

35. Mayo-Munoz, D., Pinilla-Redondo, R., Birkholz, N. & Fineran, P.C. A host of armor: Prokaryotic immune strategies against mobile genetic elements. Cell Reports 42, 112672 (2023).

36. Tesson, F. et al. A comprehensive resource for exploring antiphage defense: DefenseFinder webservice, wiki and databases. Nucleic Acids Research 53, 13 (2025).

37. Leão, P. et al. Asgard archaea defense systems and their roles in the origin of eukaryotic immunity. Nature Communications 15, 6386 (2024).

38. Liu, Y., Botelho, J. & Iranzo, J. Timescales and genetic linkage explain the variable impact of defense systems on horizontal gene transfer Genome Research 35, 268–278 (2025).

39. Kogay, R., Wolf, Y.I. & Koonin, E.V. Defence systems and horizontal gene transfer in bacteria. Environmental Microbiology 26 (2024).

40. Li, M. et al. Toxin-antitoxin RNA pairs safeguard CRISPR-Cas systems. Science 372, eabe5601 (2021).

41. Dupuis, M.È., Villion, M., Magadán, A.H. & Moineau, S. CRISPR-Cas and restriction– modification systems are compatible and increase phage resistance. Nature Communications 4, 2087 (2013).

42. Wu, Y. et al. Bacterial defense systems exhibit synergistic anti-phage activity. Cell Host & Microbe 32, 557–572 (2024).

43. Maguin, P., Varble, A., Modell, J.W. & Marraffini, L.A. Cleavage of viral DNA by restriction endonucleases stimulates the type II CRISPR-Cas immune response. Molecular Cell 82, 907–919.e907 (2022).

44. Dong, X. et al. Evolutionary ecology of microbial populations inhabiting deep sea sediments associated with cold seeps. Nature Communications 14, 1127 (2023).

45. Dong, X. et al. A vast repertoire of secondary metabolites potentially influences community dynamics and biogeochemical processes in cold seeps. Science Advances 10, eadl2281 (2024).

46. Kieft, K., Zhou, Z. & Anantharaman, K. VIBRANT: automated recovery, annotation and curation of microbial viruses, and evaluation of viral community function from genomic sequences. Microbiome 8, 1–23 (2020).

47. Guo, J. et al. VirSorter2: a multi-classifier, expert-guided approach to detect diverse DNA and RNA viruses. Microbiome 9, 1–13 (2021).

48. Camargo, A.P. et al. Identification of mobile genetic elements with geNomad. Nature Biotechnology 42, 1303–1312 (2023).

49. Roux, S. et al. Minimum Information about an Uncultivated Virus Genome (MIUViG). Nature Biotechnology 37, 29–37 (2019).

50. Roux, S. et al. iPHoP: An integrated machine learning framework to maximize host prediction for metagenome-derived viruses of archaea and bacteria. PLoS Biology 21, e3002083 (2023).

51. Zhou, K. et al. Genomic and transcriptomic insights into complex virus-prokaryote interactions in marine biofilms. The ISME Journal 17, 2303–2312 (2023).

52. Silas, S. et al. Activation of bacterial programmed cell death by phage inhibitors of host immunity. Molecular Cell 85, 1838–1851.e10 (2025).

53. Yirmiya, E. et al. Phages overcome bacterial immunity via diverse anti-defence proteins. Nature 625, 352–359 (2024).

54. Yang, L. et al. Insights into the inhibition of type I-F CRISPR-Cas system by a multifunctional anti-CRISPR protein AcrIF24. Nature Communications 13, 1931 (2022).

55. Suresh, S.K., Murugan, K. & Sashital, D.G. Enzymatic anti-CRISPRs improve the bacteriophage arsenal. Nature Structural & Molecular Biology 26, 250–251 (2019).

56. Knott, G.J. et al. Broad-spectrum enzymatic inhibition of CRISPR-Cas12a. Nature Structural & Molecular Biology 26, 315–321 (2019).

57. Müller, A.L., Kjeldsen, K.U., Rattei, T., Pester, M. & Loy, A. Phylogenetic and environmental diversity of DsrAB-type dissimilatory (bi)sulfite reductases. The ISME Journal 9, 1152–1165 (2015).

58. Gu, Y. et al. Bacterial Shedu immune nucleases share a common enzymatic core regulated by diverse sensor domains. Molecular Cell 85, 523–536 (2024).

59. Dong, X. et al. Phylogenetically and catabolically diverse diazotrophs reside in deep-sea cold seep sediments. Nature Communications 13, 4885 (2022).

60. Sheng, Y. et al. Insertion sequence transposition inactivates CRISPR-Cas immunity. Nature Communications 14, 4366 (2023).

61. Koonin, E.V., Makarova, K.S. & Zhang, F. Diversity, classification and evolution of CRISPR-Cas systems. Current Opinion in Microbiology 37, 67–78 (2017).

62. Martínez Arbas, S., et al. Roles of bacteriophages, plasmids and CRISPR immunity in microbial community dynamics revealed using time-series integrated meta-omics. Nature Microbiology 6, 123–135 (2021).

63. Sun, C.L., Thomas, B.C., Barrangou, R. & Banfield, J.F. Metagenomic reconstructions of bacterial CRISPR loci constrain population histories. The ISME Journal 10, 858–870 (2016).

64. Guerrero, L.D. et al. Long-run bacteria-phage coexistence dynamics under natural habitat conditions in an environmental biotechnology system. The ISME Journal 15, 636–648 (2021).

65. Huo, Y. et al. Structural and biochemical insights into the mechanism of the Gabija bacterial immunity system. Nature Communications 15, 836 (2024).

66. Olm, M.R., Brown, C.T., Brooks, B. & Banfield, J.F. dRep: a tool for fast and accurate genomic comparisons that enables improved genome recovery from metagenomes through de-replication. The ISME journal 11, 2864–2868 (2017).

67. Parks, D.H. et al. GTDB: an ongoing census of bacterial and archaeal diversity through a phylogenetically consistent, rank normalized and complete genome-based taxonomy. Nucleic Acids Research 50, D785–D794 (2022).

68. Nguyen, L.T., Schmidt, H.A., Von Haeseler, A. & Minh, B.Q. IQ-TREE: a fast and effective stochastic algorithm for estimating maximum-likelihood phylogenies. Molecular Biology and Evolution 32, 268–274 (2015).

69. Letunic, I. & Bork, P. Interactive Tree Of Life (iTOL) v5: an online tool for phylogenetic tree display and annotation. Nucleic Acids Research 49, W293–W296 (2021).

70. Breitwieser, F.P. & Salzberg, S.L. Pavian: interactive analysis of metagenomics data for microbiome studies and pathogen identification. Bioinformatics 36, 1303–1304 (2020).

71. Hyatt, D. et al. Prodigal: prokaryotic gene recognition and translation initiation site identification. BMC Bioinformatics 11, 1–11 (2010).

72. Zhang, C. et al. The majority of microorganisms in gas hydrate-bearing subseafloor sediments ferment macromolecules. Microbiome 11, 37 (2023).

73. Xiao, X. et al. Metal-driven anaerobic oxidation of methane as an important methane sink in methanic cold seep sediments. Microbiology Spectrum 11, e05337–05322 (2023).

74. Li, W.L. et al. Microbial ecology of sulfur cycling near the sulfate–methane transition of deeplJsea cold seep sediments. Environmental Microbiology 23, 6844–6858 (2021).

75. Uritskiy, G.V., DiRuggiero, J. & Taylor, J. MetaWRAP—a flexible pipeline for genome-resolved metagenomic data analysis. Microbiome 6, 1–13 (2018).

76. Kopylova, E., Noé, L. & Touzet, H. SortMeRNA: fast and accurate filtering of ribosomal RNAs in metatranscriptomic data. Bioinformatics 28, 3211–3217 (2012).

77. Patro, R., Duggal, G., Love, M.I., Irizarry, R.A. & Kingsford, C. Salmon provides fast and bias-aware quantification of transcript expression. Nature Methods 14, 417–419 (2017).

78. Nayfach, S. et al. CheckV assesses the quality and completeness of metagenome-assembled viral genomes. Nature Biotechnology 39, 578–585 (2021).

79. Shaffer, M. et al. DRAM for distilling microbial metabolism to automate the curation of microbiome function. Nucleic Acids Research 48, 8883–8900 (2020).

80. Ye, J., McGinnis, S. & Madden, T.L. BLAST: improvements for better sequence analysis. Nucleic Acids Research 34, W6–W9 (2006).

81. 81. Coutinho, F.H., et al. RaFAH: Host prediction for viruses of Bacteria and Archaea based on protein content. Patterns 2, 100274 (2021).

82. Galiez, C., Siebert, M., Enault, F., Vincent, J. & Söding, J. WIsH: who is the host? Predicting prokaryotic hosts from metagenomic phage contigs. Bioinformatics 33, 3113–3114 (2017).

83. 83. Wang, W., et al. A network-based integrated framework for predicting virus–prokaryote interactions. NAR Genomics and Bioinformatics 2, lqaa044 (2020).

84. Lu, C. et al. Prokaryotic virus host predictor: a Gaussian model for host prediction of prokaryotic viruses in metagenomics. BMC Biology 19, 5 (2021).

85. Couvin, D. et al. CRISPRCasFinder, an update of CRISRFinder, includes a portable version, enhanced performance and integrates search for Cas proteins. Nucleic Acids Research 46, W246–W251 (2018).

86. Grandjean, M. Gephi: Introduction to network analysis and visualisation. International AAAI Conference on Weblogs and Social Media (2015).

87. Yan, Y., Zheng, J., Zhang, X. & Yin, Y. dbAPIS: a database of anti-prokaryotic immune system genes. Nucleic Acids Research 52, 419–425 (2023).

88. Buchfink, B., Reuter, K. & Drost, H.G. Sensitive protein alignments at tree-of-life scale using DIAMOND. Nature Methods 18, 366–368 (2021).

89. Mazzocco, A., Waddell, T.E., Lingohr, E. & Johnson, R.P. Enumeration of bacteriophages using the small drop plaque assay system. Bacteriophages: methods and protocols, volume 1: isolation, characterization, and interactions, 81–85 (2009).

90. Khedkar, S. et al. Landscape of mobile genetic elements and their antibiotic resistance cargo in prokaryotic genomes. Nucleic Acids Research 50, 3155–3168 (2022).

91. Neukirchen, S. & Sousa, F.L. DiSCo: a sequence-based type-specific predictor of Dsr-dependent dissimilatory sulphur metabolism in microbial data. Microbial Genomics 7, 000603 (2021).

92. Zhou, Z. et al. METABOLIC: high-throughput profiling of microbial genomes for functional traits, metabolism, biogeochemistry, and community-scale functional networks. Microbiome 10, 33 (2022).

93. Tu, Q., Lin, L., Cheng, L., Deng, Y. & He, Z. NCycDB: a curated integrative database for fast and accurate metagenomic profiling of nitrogen cycling genes. Bioinformatics 35, 1040–1048 (2019).

94. Nakamura, T., Yamada, K.D., Tomii, K. & Katoh, K. Parallelization of MAFFT for large-scale multiple sequence alignments. Bioinformatics 34, 2490–2492 (2018).

95. Capella-Gutiérrez, S., Silla-Martínez, J.M. & Gabaldón, T. trimAl: a tool for automated alignment trimming in large-scale phylogenetic analyses. Bioinformatics 25, 1972–1973 (2009).

96. Chen, C. et al. TBtools-II: A “one for all, all for one” bioinformatics platform for biological big-data mining. Molecular Plant 16, 1733–1742 (2023).

97. Paysan-Lafosse, T., et al. InterPro in 2022. Nucleic Acids Research 51, D418–D427 (2023).

98. Waterhouse, A.M., Procter, J.B., Martin, D.M.A., Clamp, M. & Barton, G.J. Jalview Version 2—a multiple sequence alignment editor and analysis workbench. Bioinformatics 25, 1189–1191 (2009).

99. Mirdita, M., Steinegger, M. & Söding, J. MMseqs2 desktop and local web server app for fast, interactive sequence searches. Bioinformatics 35, 2856–2858 (2019).

100. Mirdita, M. et al. ColabFold: making protein folding accessible to all. Nature Methods 19, 679–682 (2022).

101. Jumper, J. et al. Highly accurate protein structure prediction with AlphaFold. nature 596, 583–589 (2021).

102. Abramson, J. et al. Accurate structure prediction of biomolecular interactions with AlphaFold 3. Nature 630, 493–500 (2024).

103. Van Kempen, M. et al. Fast and accurate protein structure search with Foldseek. Nature biotechnology 42, 243–246 (2024).

104. DeLano, W.L. Pymol: An open-source molecular graphics tool. CCP4 Newsl. Protein Crystallogr 40, 82–92 (2002).

