## Supplementary for "Co-occurrence of diverse defense systems shapes complex microbe-virus relationships in deep-sea cold seeps": Supplementary Information.docx

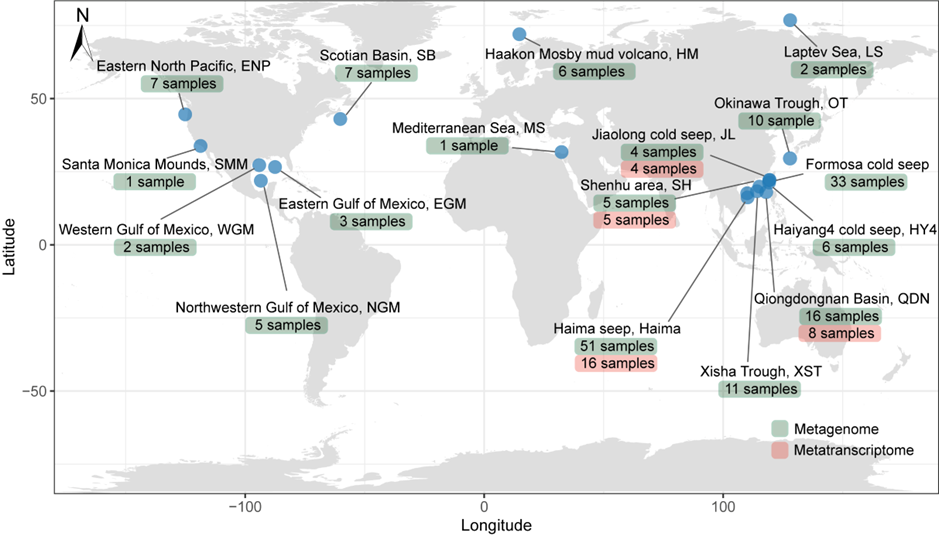

**Supplementary** **Figure 1. Geographic locations of cold seep study sites**. The map indicates the distribution of cold seep sites where metagenomic and metatranscriptomic data were collected. Sites marked with green squares represent locations where metagenomic data were obtained, while sites with orange squares indicate metatranscriptomic data collection. The world map was generated using the ggplot2 package in R version 4.3.3, modified from Han et al. (2023).

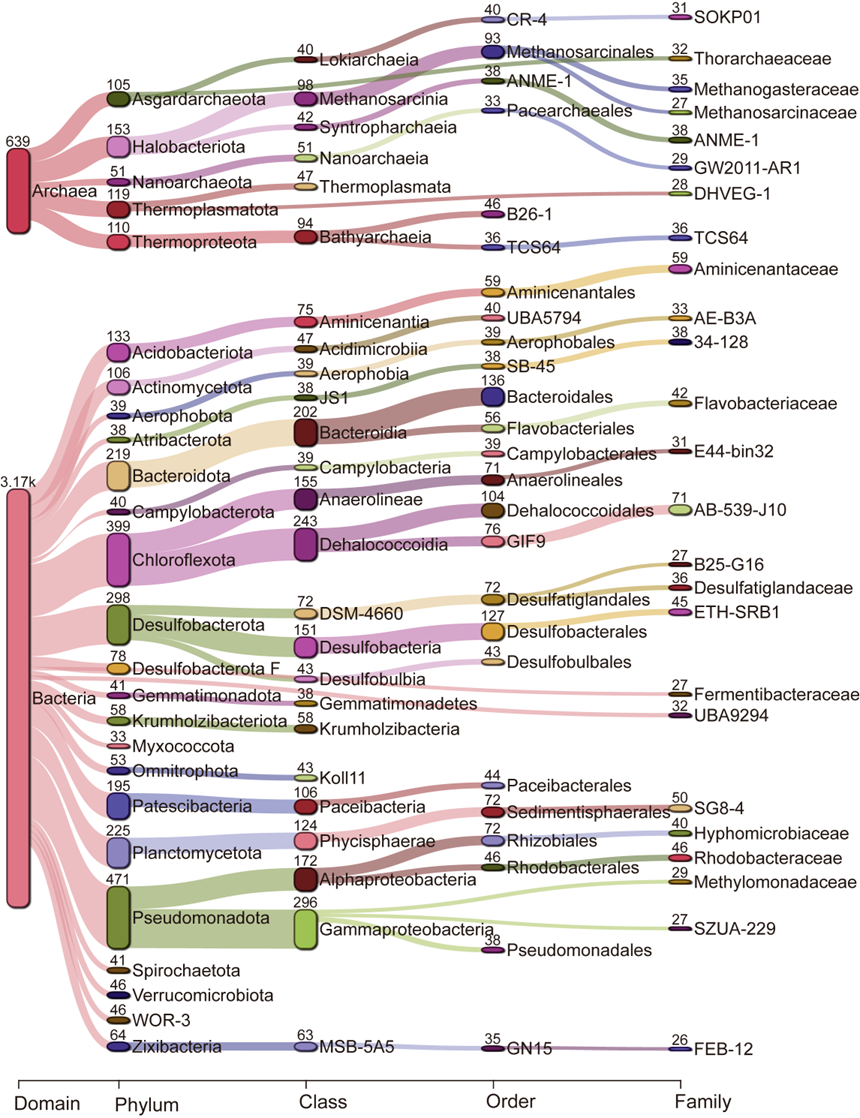

**Supplementary Figure 2. Sankey diagram illustrating the taxonomic classification of MAGs from cold seep sediment prokaryotes.** The distribution of archaeal and bacterial communities is depicted across taxonomic levels from domain to family. Numbers indicate the quantity of recovered MAGs at each taxonomic level. Detailed species classification information is provided in **Supplementary Data 3**.

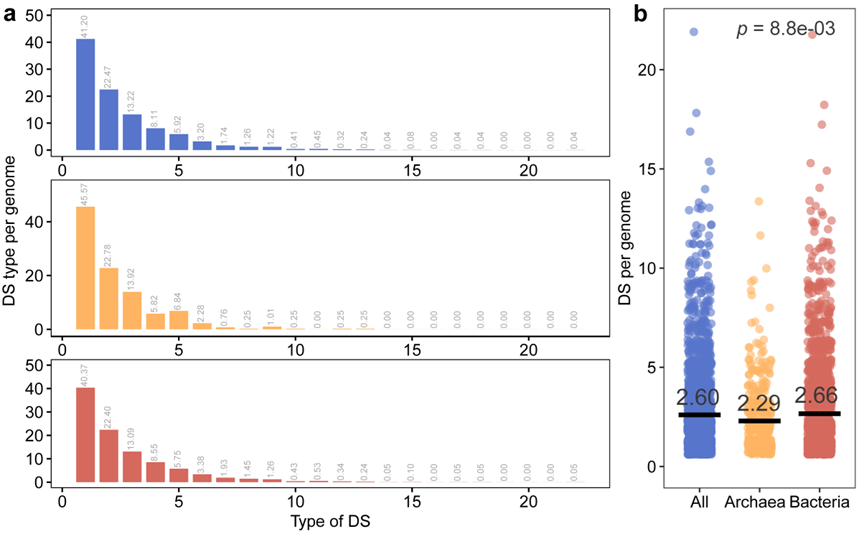

**Supplementary Figure 3. Number of distinct defense system (DS) types per MAG.** Blue bars represent all prokaryotes, orange bars represent archaea, and red bars represent bacteria. **(a)** Distribution of the number of defense systems (DS) types per MAG for all prokaryotes (blue, n=2,466), archaea (orange, n=395), and bacteria (red, n= 2,071). **(b)** Number of defense systems (DSs) types per MAG for all prokaryotes (blue, n=2,466), archaea (orange, n=395), and bacteria (red, n= 2,071), with black lines indicating the average number. Statistical differences between archaea and bacteria were assessed using a two-sided Wilcoxon rank-sum test (*p*=8.8e-03). Detailed statistical data are provided in **Supplementary Data 5**.

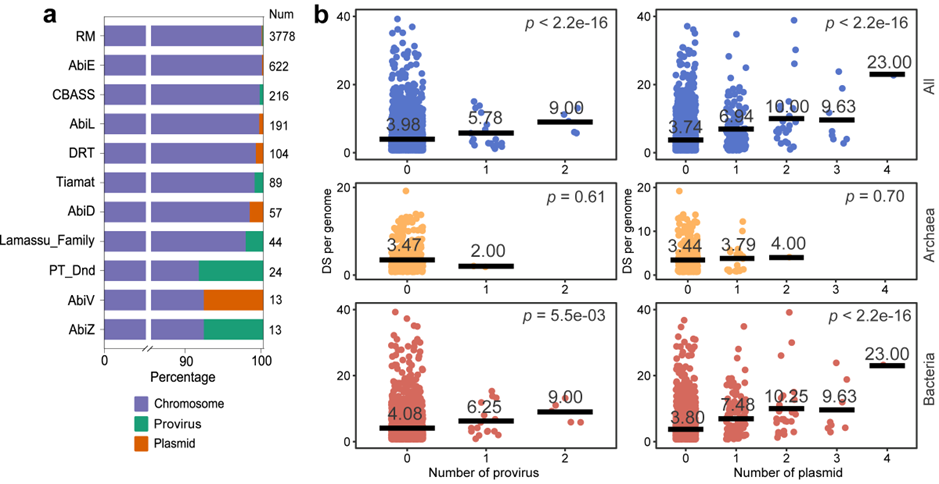

**Supplementary Figure 4. Association between mobile genetic elements (MGEs) and the distribution of defense systems in cold seep prokaryotes. (a)** Distribution and proportion of 11 defense systems found on MGEs. **(b)** Effect of the number of prophages and plasmids within MAGs on the quantity of defense systems for all prokaryotes (blue, n=2,466), archaea (orange, n=395), and bacteria (red, n=2,071). Black lines and numbers denote the corresponding average number of defense systems. *p* value is calculated using a two-sided Pearson correlation test to assess the linear relationship between the number of defense systems and MGEs.

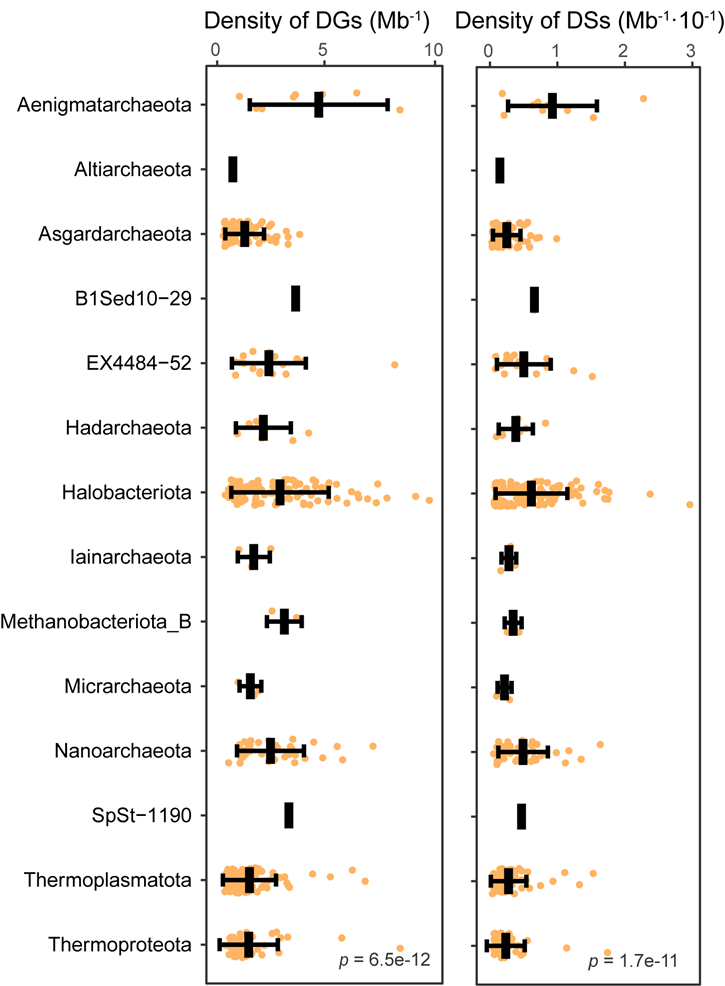

**Supplementary Figure 5. Density of defense systems and defense genes in archaea.** The density of defense systems (per Mb) and defense genes (per Mb × 10⁻¹) in archaeal MAGs are shown across different phyla. Statistical differences among phyla were assessed using a two-sided Kruskal-Wallis rank-sum test. Data are presented as mean ± SD.

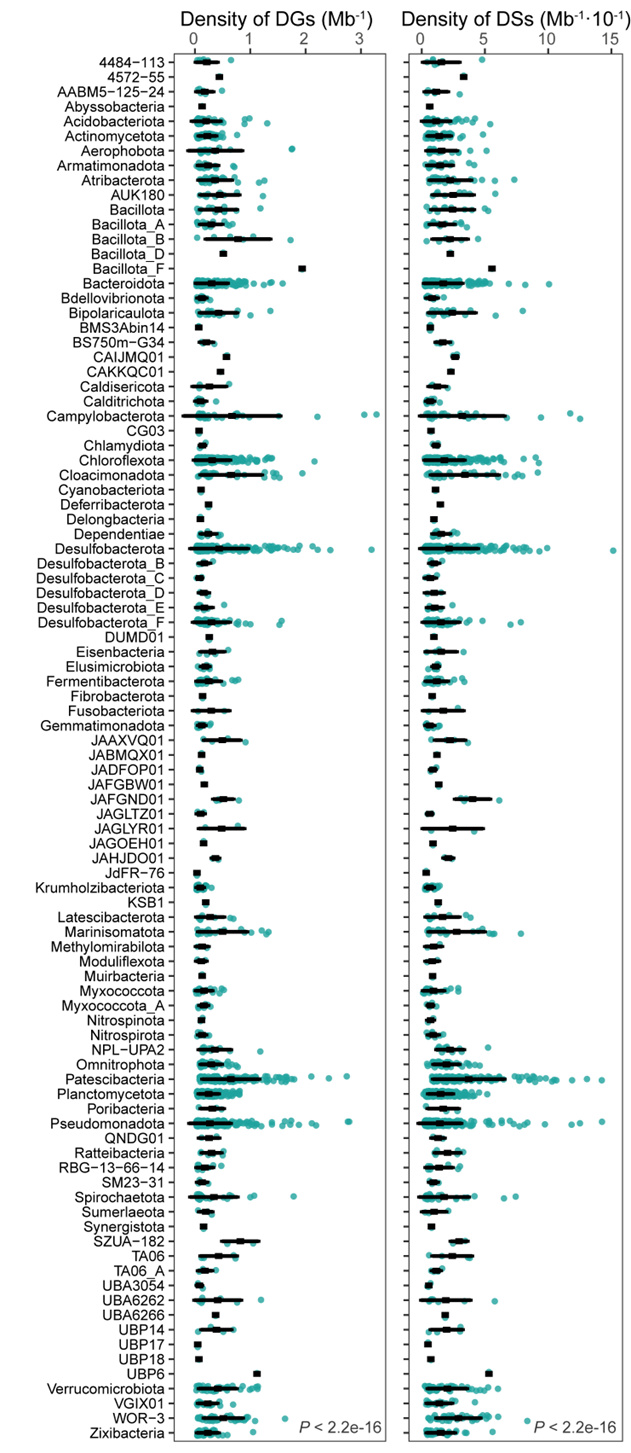

**Supplementary Figure 6. Density of defense systems and defense genes in bacteria.** The density of defense systems (per Mb) and defense genes (per Mb × 10⁻¹) in bacterial MAGs are shown across different phyla. Statistical differences among phyla were assessed using a two-sided Kruskal-Wallis rank-sum test. Data are presented as mean ± SD.

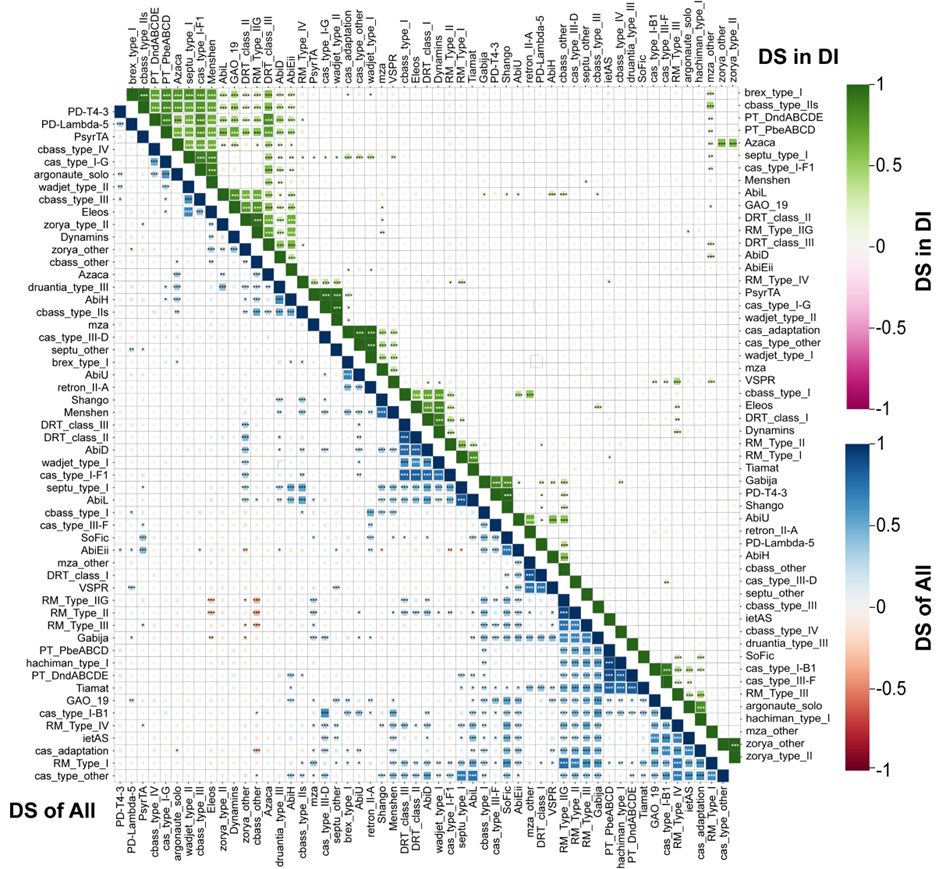

**Supplementary Figure 7. Co-occurrence patterns of defense subsystems in cold seep prokaryotes.** Correlations among 54 defense subsystems in cold seep prokaryotes. The lower-left matrix (blue-red gradient) displays correlations among all defense subsystems across prokaryotic MAGs, while the upper-right matrix (green-pink gradient) shows correlations within defense islands. Pearson correlation coefficients were calculated based on system abundance using two-sided tests; coefficients are color-coded, and larger squares indicate stronger correlations. Significance levels are denoted by asterisks:* *p* ≤ 0.05, ** *p* ≤ 0.01, *** *p* ≤ 0.001. Detailed data are provided in **Supplementary Data 8**.

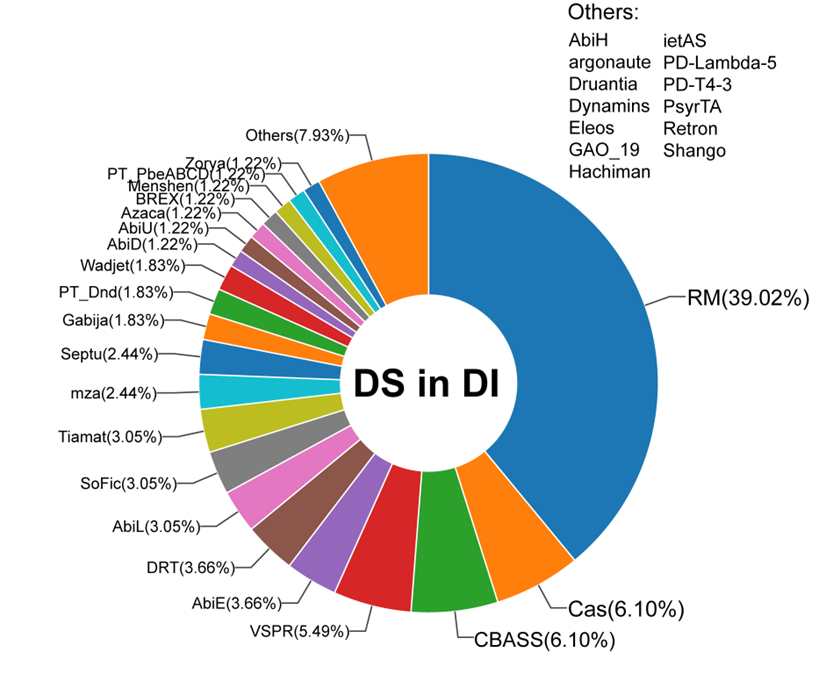

**Supplementary Figure 8. Composition of defense systems within defense islands.** The proportions of various defense system types are displayed that are present on defense islands.

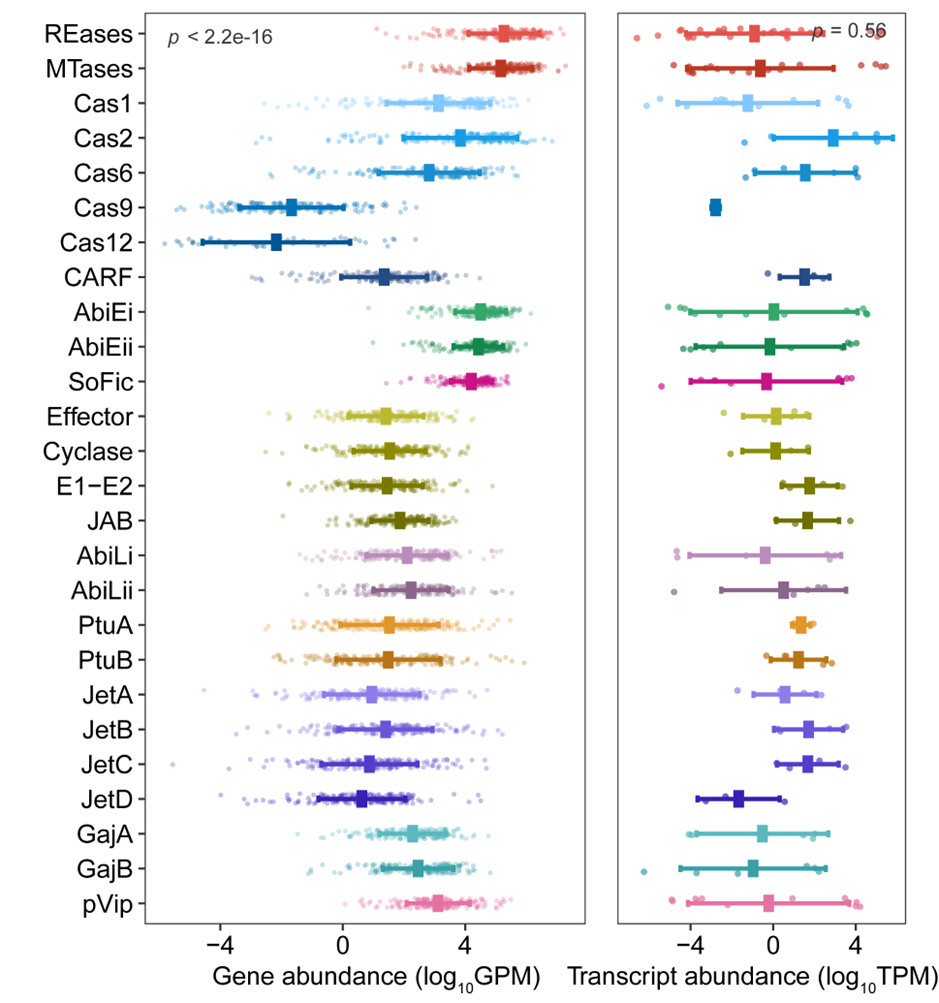

**Supplementary Figure 9. Gene abundance and expression levels of defense genes in cold seep prokaryotes.** Gene abundance (genes per million, GPM) and transcript abundance (transcripts per million, TPM) are shown for 26 genes corresponding to the top ten defense systems across different sediment samples. Different colors represent different types of defense systems. Statistical differences among defense system types or genes were assessed using a two-sided Kruskal-Wallis rank-sum test. Data are presented as mean ± SD.

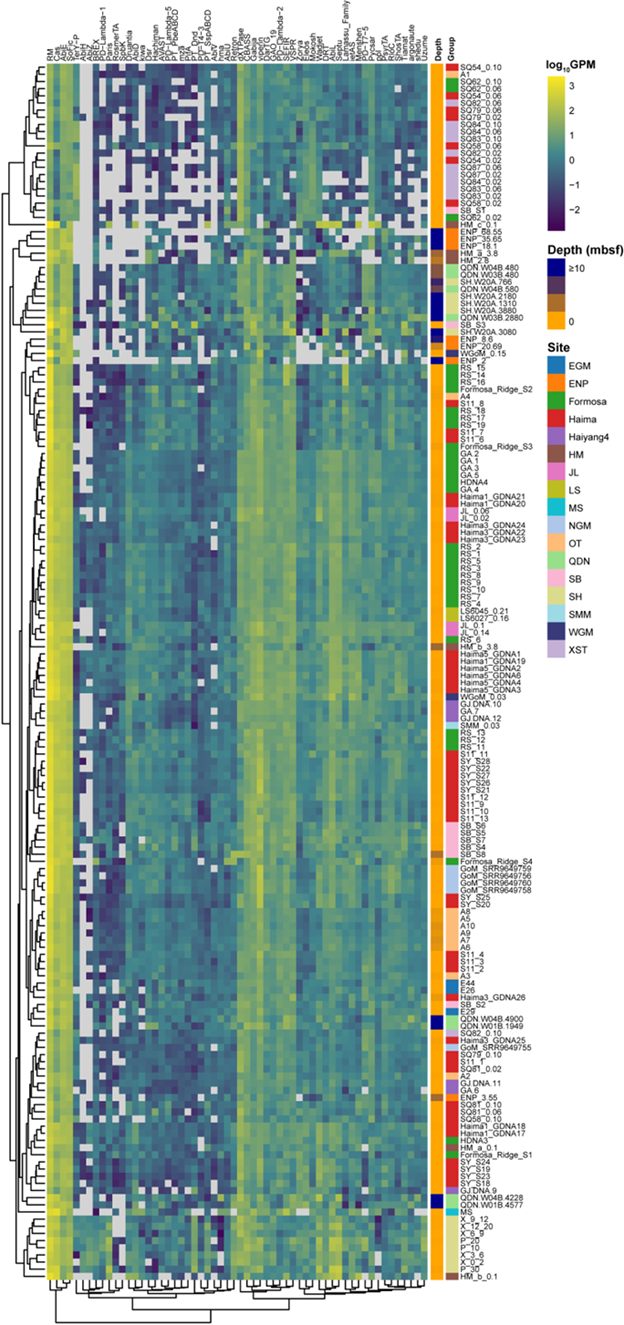

**Supplementary Figure 10. Gene abundance of various defense systems across all cold seep samples.** Gene abundance was measured as genes per million (GPM). Gray squares indicate that the gene abundance for that system is zero in the respective sample. Sample depths and locations are indicated on the right.

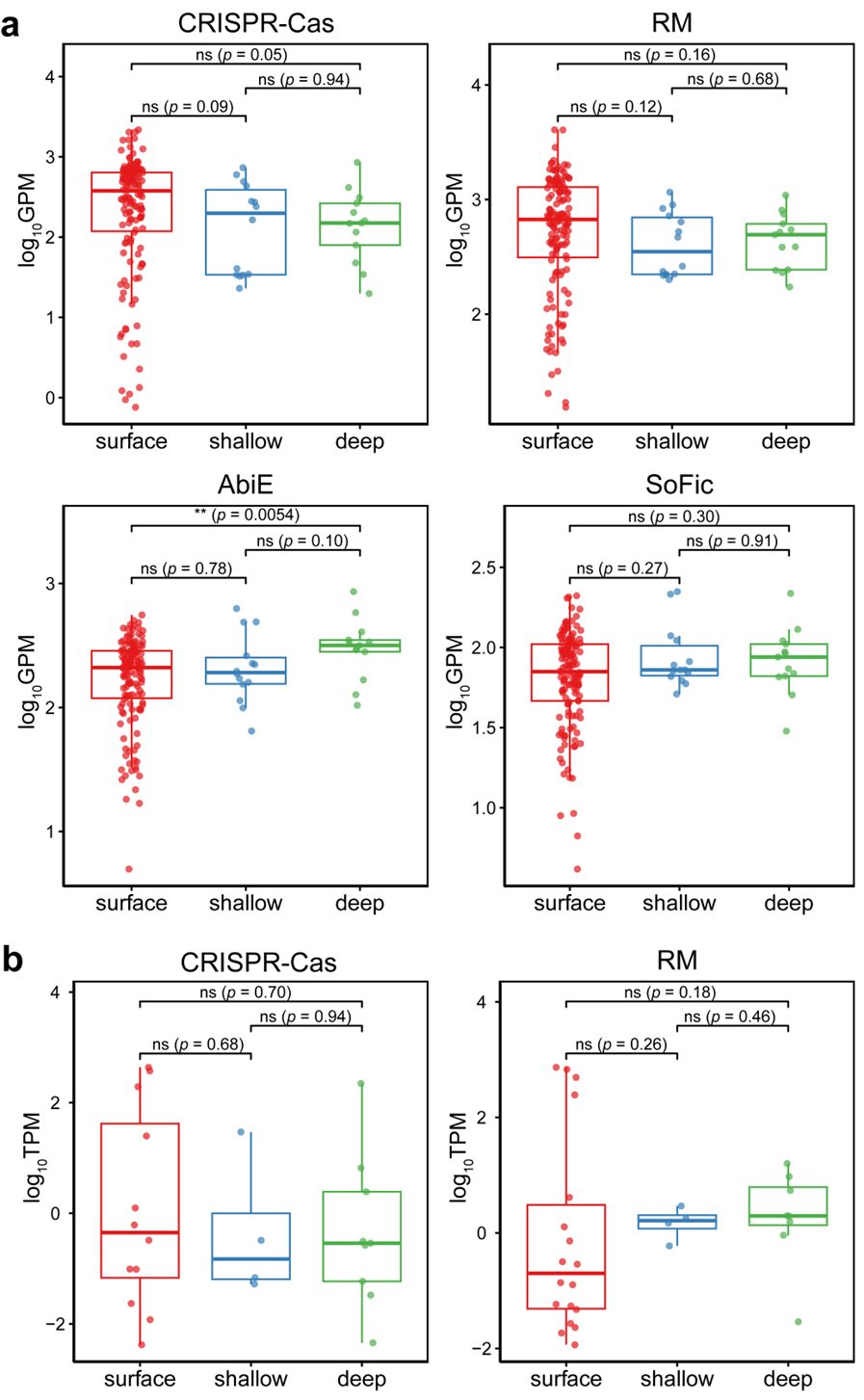

**Supplementary Figure 11. Gene abundance (a) and expression levels (b) of four major types of defense systems at different sediment depths.** Gene abundance was measured as genes per million (GPM) and transcript abundance as transcripts per million (TPM). AbiE and SoFic systems were rarely detected and are therefore not shown. Sediment depths are categorized into surface (<1 meters below seafloor [mbsf]), shallow (1–10 mbsf), and deep (>10 mbsf) groups. Differences were computed using a two-sided Wilcoxon rank-sum test. Boxplot: center line, median; box limits, upper and lower quartiles; whiskers, 1.5 × interquartile range; points, individual data values.

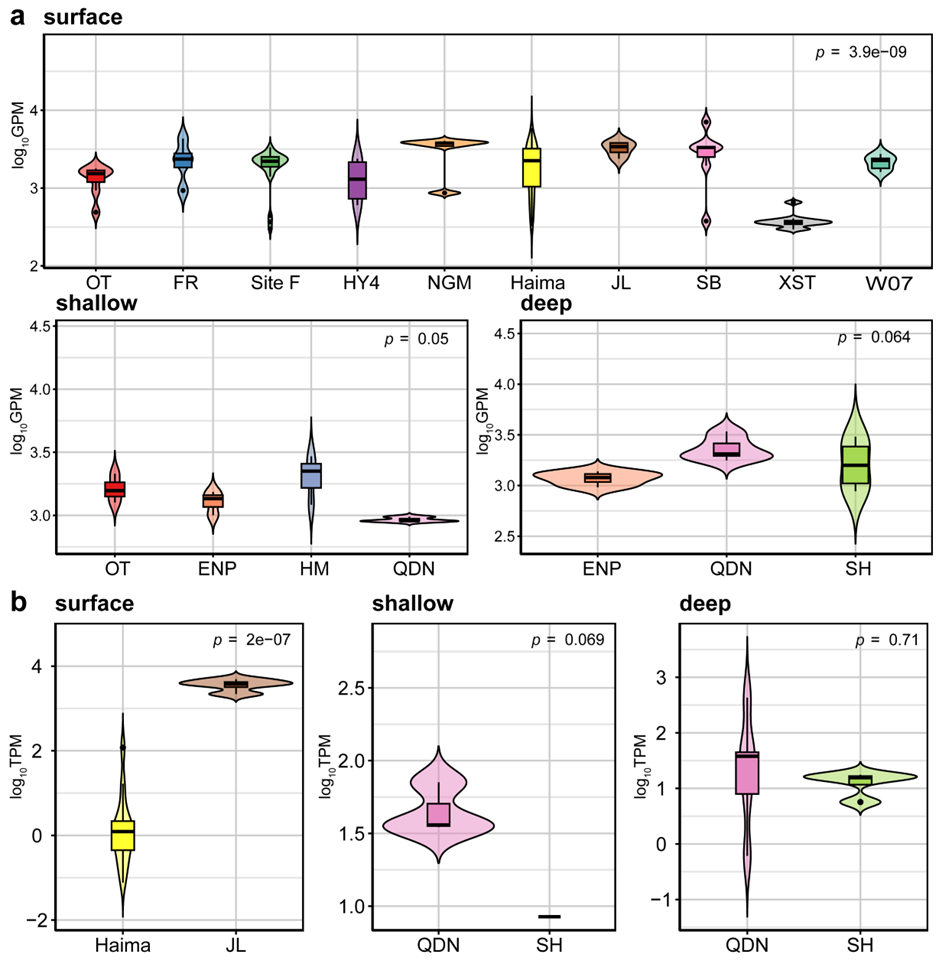

**Supplementary Figure 12. Differences in defense system gene abundance (a) and transcript abundance (b) at different depths of cold seeps from various sites.** Gene abundance was measured as genes per million (GPM) and transcript abundance as transcripts per million (TPM). Sediment depths are categorized into three groups: surface (<1 mbsf), shallow (1–10 mbsf), and deep (>10 mbsf). Differences were computed using a two-sided Wilcoxon rank-sum test. Boxplot: center line, median; box limits, upper and lower quartiles; whiskers, 1.5 × interquartile range.

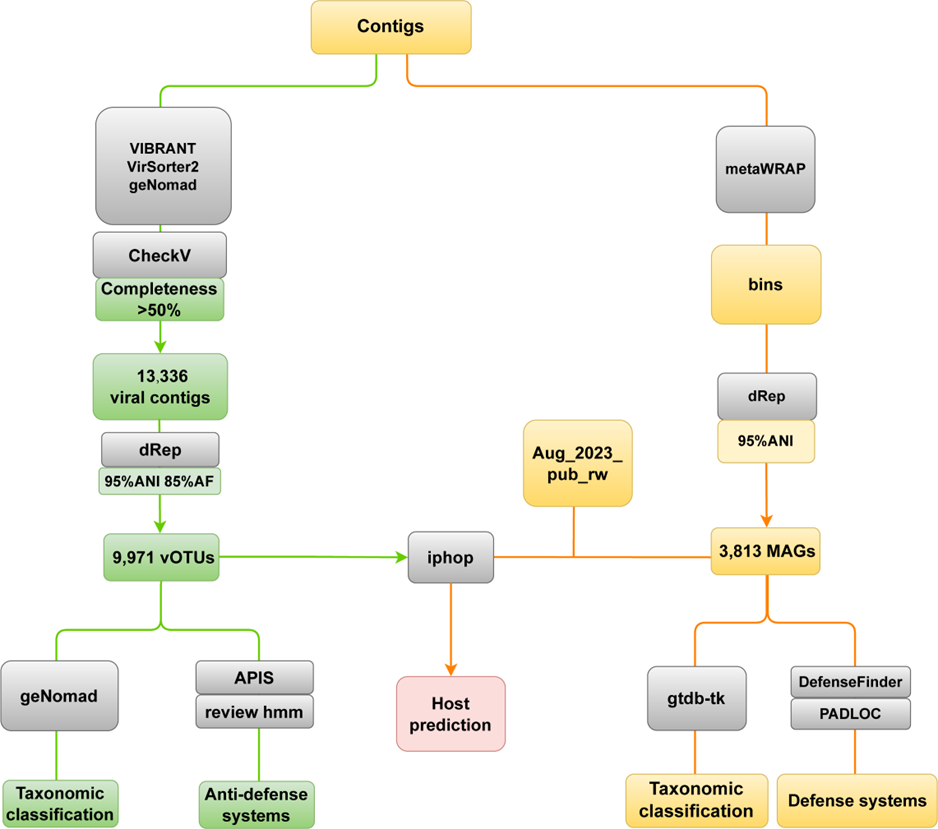

**Supplementary Figure 13. Workflow of virus-related analysis conducted in this study.** Green boxes represent the identification of cold seep viruses and viral anti-defense genes; yellow boxes denote the processing of cold seep prokaryotic MAGs and the identification of defense systems; red boxes indicate the determination of virus-host linkages.

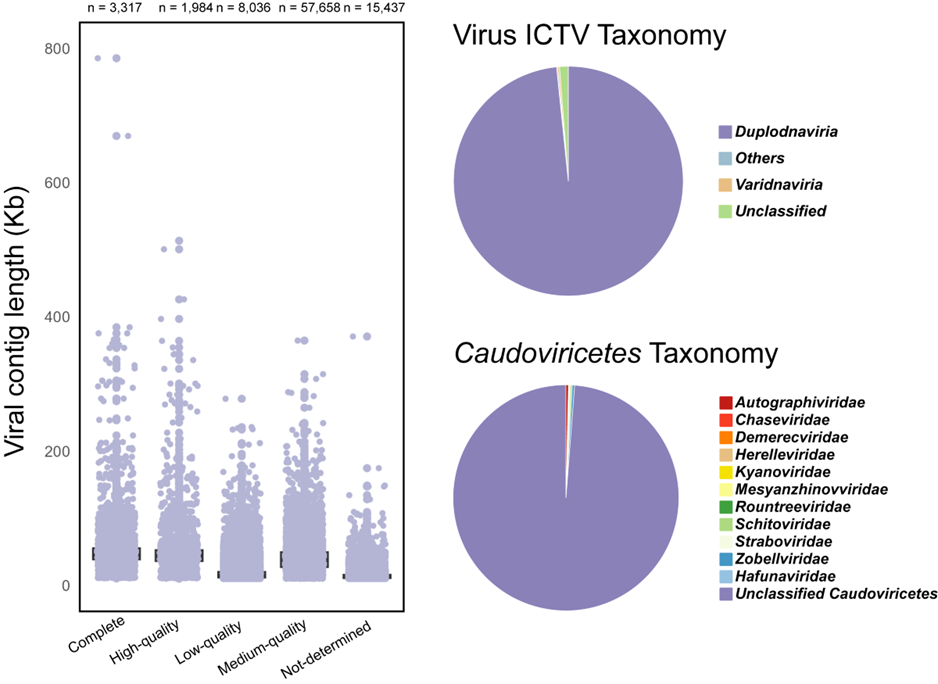

**Supplementary Figure 14. Characteristics of cold seep viruses.** Quality assessment of viral contigs was evaluated by CheckV. Viruses are classified according to the International Committee on Taxonomy of Viruses (ICTV) at the phylum level, and *Caudoviricetes* viruses are further classified at the family level.

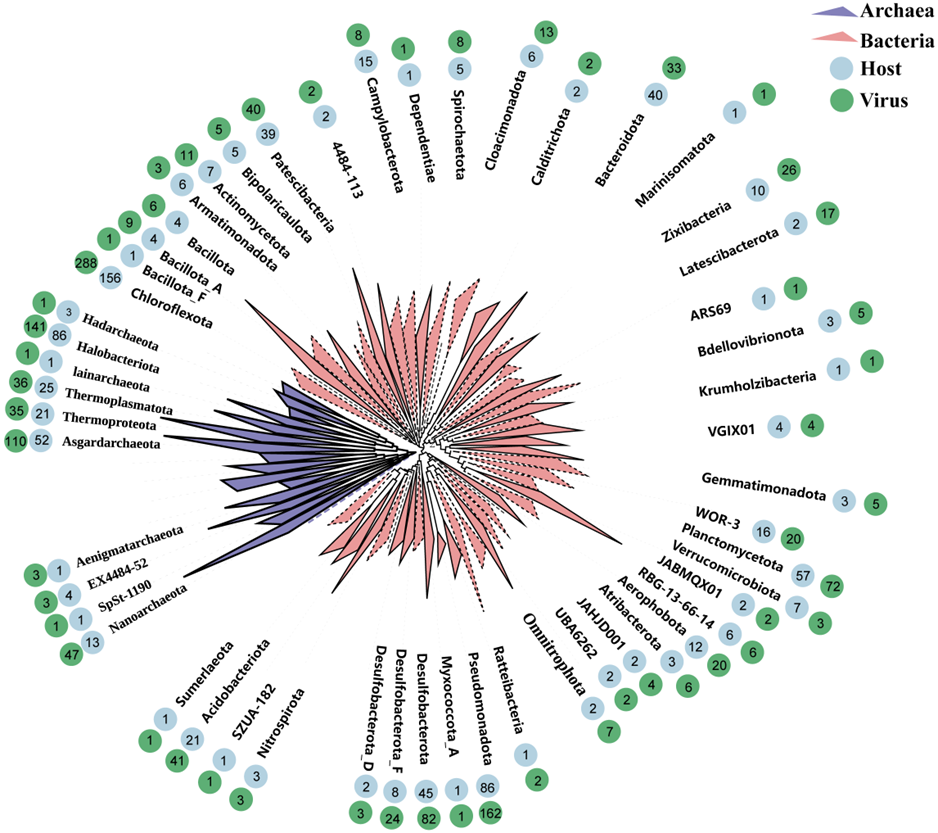

**Supplementary Figure 15. Virus-host linkages identified in cold seeps.** Maximum-likelihood phylogenetic trees of bacterial and archaeal MAGs at the phylum level were constructed based on concatenated alignments of 120 bacterial (red) or 53 archaeal (purple) single-copy marker genes. The number of host MAGs in each phylum is indicated in blue, and the number of viruses infecting hosts in each phylum is shown in green.

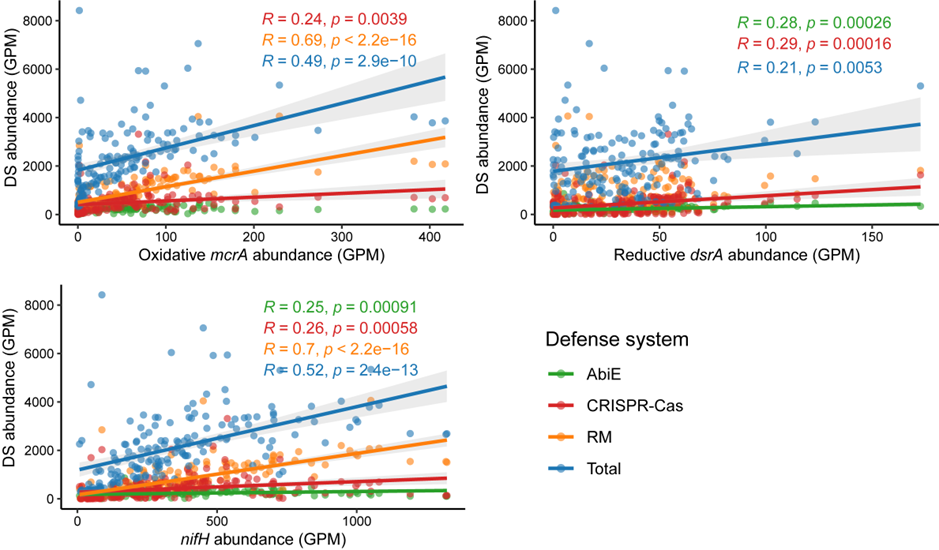

**Supplementary Figure 16. Relationship between the abundance of key metabolic genes and defense systems in cold seeps.** The abundance of RM (orange), CRISPR-Cas (red), and AbiE (green) systems, as well as the total defense system abundance (Total, blue), is plotted against the abundance of key metabolic genes including oxidative *mcrA* (anaerobic methane oxidation), reductive *dsrA* (sulfate reduction), and *nifH* (nitrogen fixation). Each data point represents the average gene abundance for a sample. Linear regression lines and *R* values are shown for each defense system type, color-coded accordingly. Correlations were assessed using a two-sided Pearson correlation test.

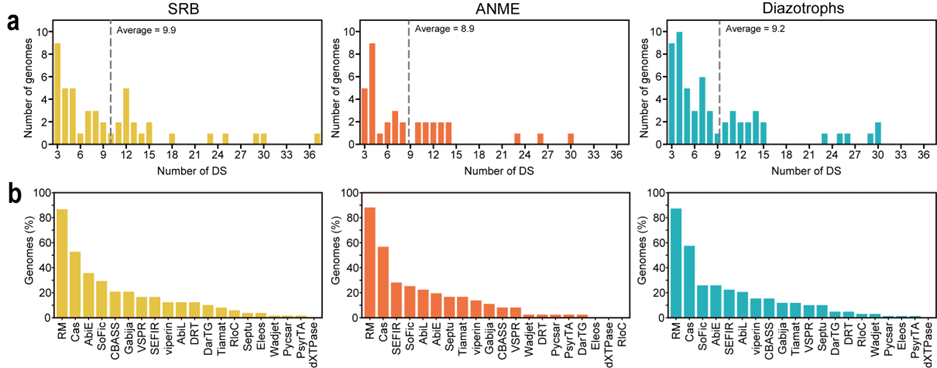

**Supplementary Figure 17.** **Distribution of defense systems in key metabolic microbial groups of cold seeps. (a)** Distribution and average number of defense system (DS) types per MAG in sulfate-reducing bacteria (SRB, yellow), anaerobic methanotrophic archaea (ANME, orange), and diazotrophs (blue). The overall average is indicated by a gray dashed line. **(b)** Proportion of MAGs within each microbial group that contain various types of defense systems.

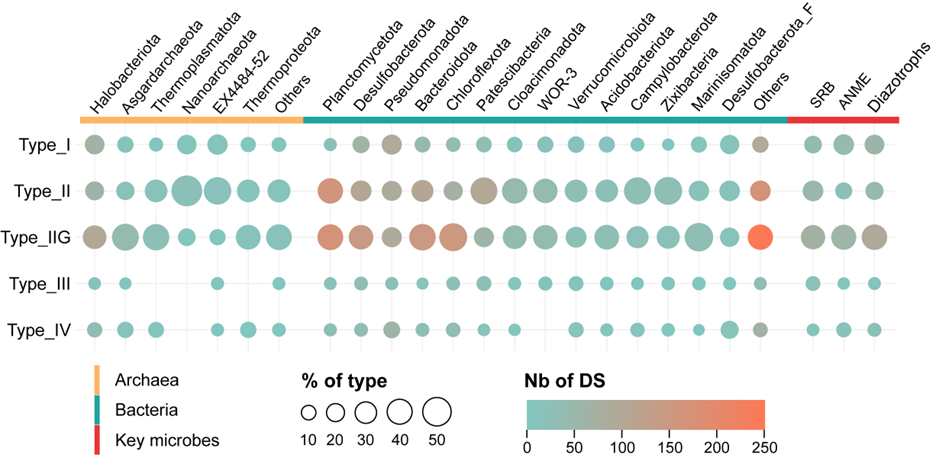

**Supplementary Figure 18.** **Distribution of restriction-modification (RM) systems among different prokaryotes in cold seeps.** The proportion and number of five types of RM systems are shown in archaea (orange), bacteria (green) at the phylum level, as well as three key microbial groups (red) in cold seeps.

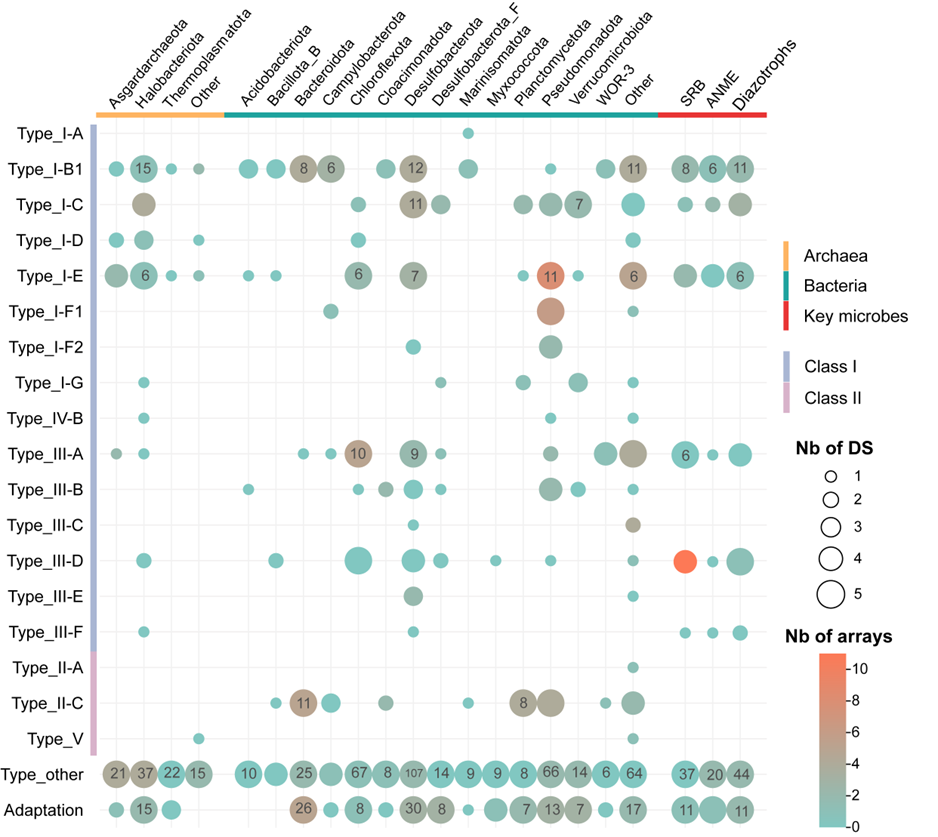

**Supplementary Figure 19. Distribution of CRISPR-Cas system among different prokaryotes in cold seeps.** The number and counts of detected arrays are displayed for 20 types of CRISPR-Cas systems (Class I: purple; Class II: pink) in archaea (orange) and bacteria (green) at the phylum level, as well as in three key microbial groups (red) in cold seeps.

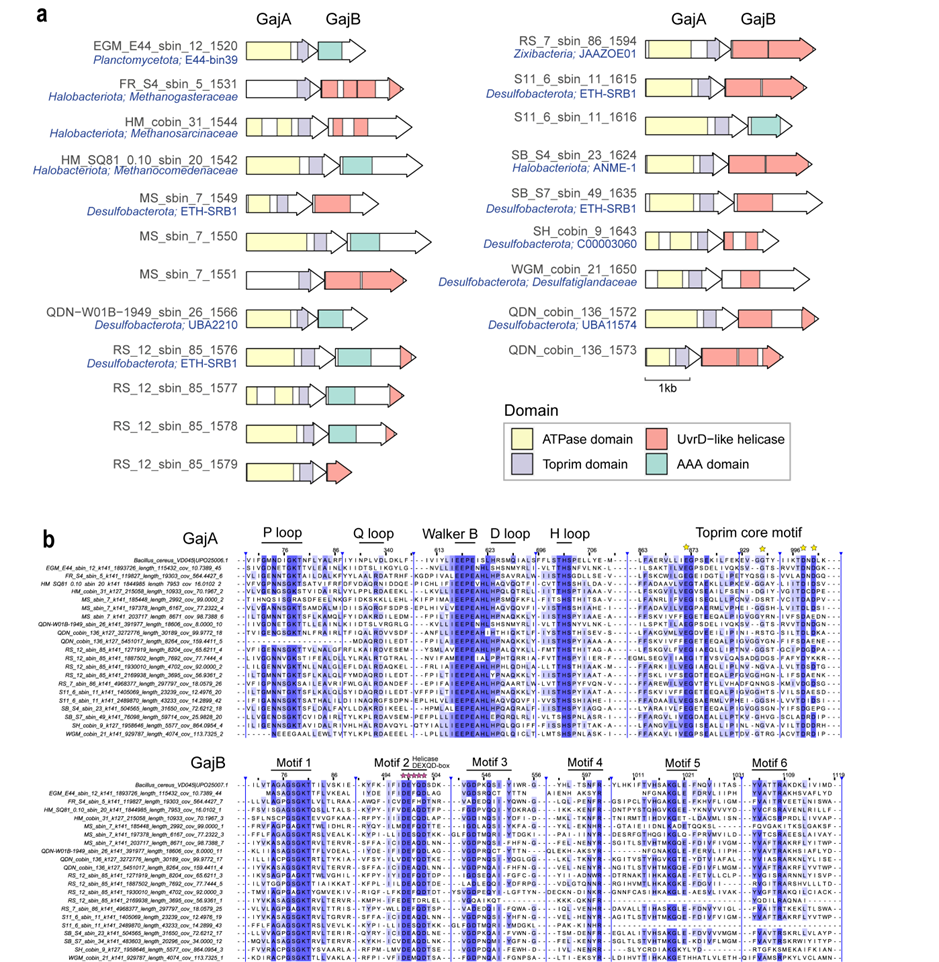

**Supplementary Figure 20. Protein domains and motifs of the Gabija system in key metabolic microbial groups. (a)** Domain organization of the GajA and GajB proteins of the Gabija system in 21 pairs of key cold seep prokaryotes. **(b)** Sequence alignment of GajA and GajB proteins according to amino acid conservation, with colors indicating the degree of conservation. GajA and GajB proteins from *Bacillus cereus* VD045 serve as the reference.
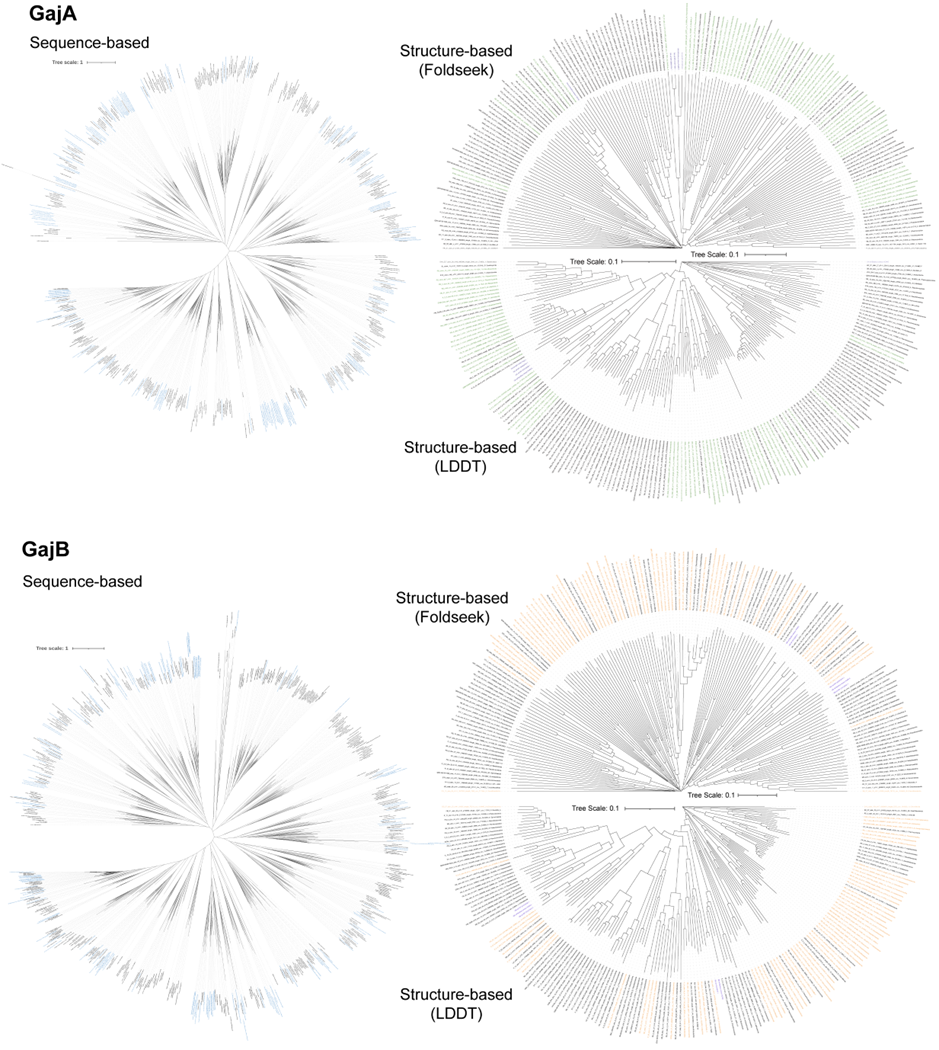

**Supplementary Figure 21.** **Phylogenetic trees of GajA and GajB proteins based on sequence and structural analyses.** In the sequence-based phylogenetic tree, sequences identified from cold seeps are shown in blue, while reference sequences are shown in black. The structural phylogenetic tree comprises two types based on Foldseek (upper semicircle) and LDDT scores (lower semicircle). Sequences containing complete domains and motifs are highlighted in green and orange.

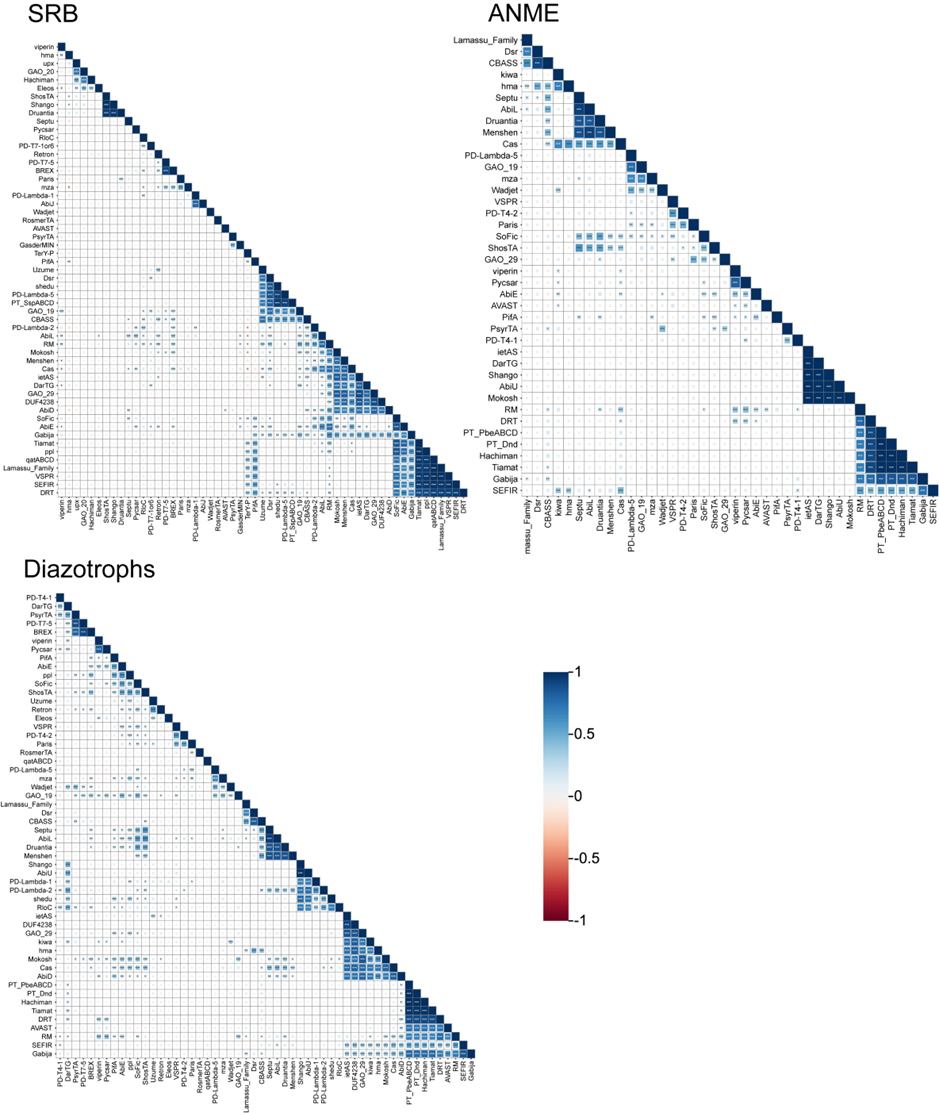

**Supplementary Figure 22.** **Correlations among defense systems in three key metabolic microbial groups in cold seeps.** Correlations between various defense systems in sulfate-reducing bacteria (SRB), anaerobic methanotrophic archaea (ANME), and diazotrophs were calculated based on system abundance using two-sided Pearson correlation coefficients. Correlation coefficients are color-coded, with larger squares indicating stronger correlations. Significance levels are denoted by asterisks: * *p* ≤ 0.05, ** *p* ≤ 0.01, *** *p* ≤ 0.001. Detailed data are provided in **Supplementary Data 16**.

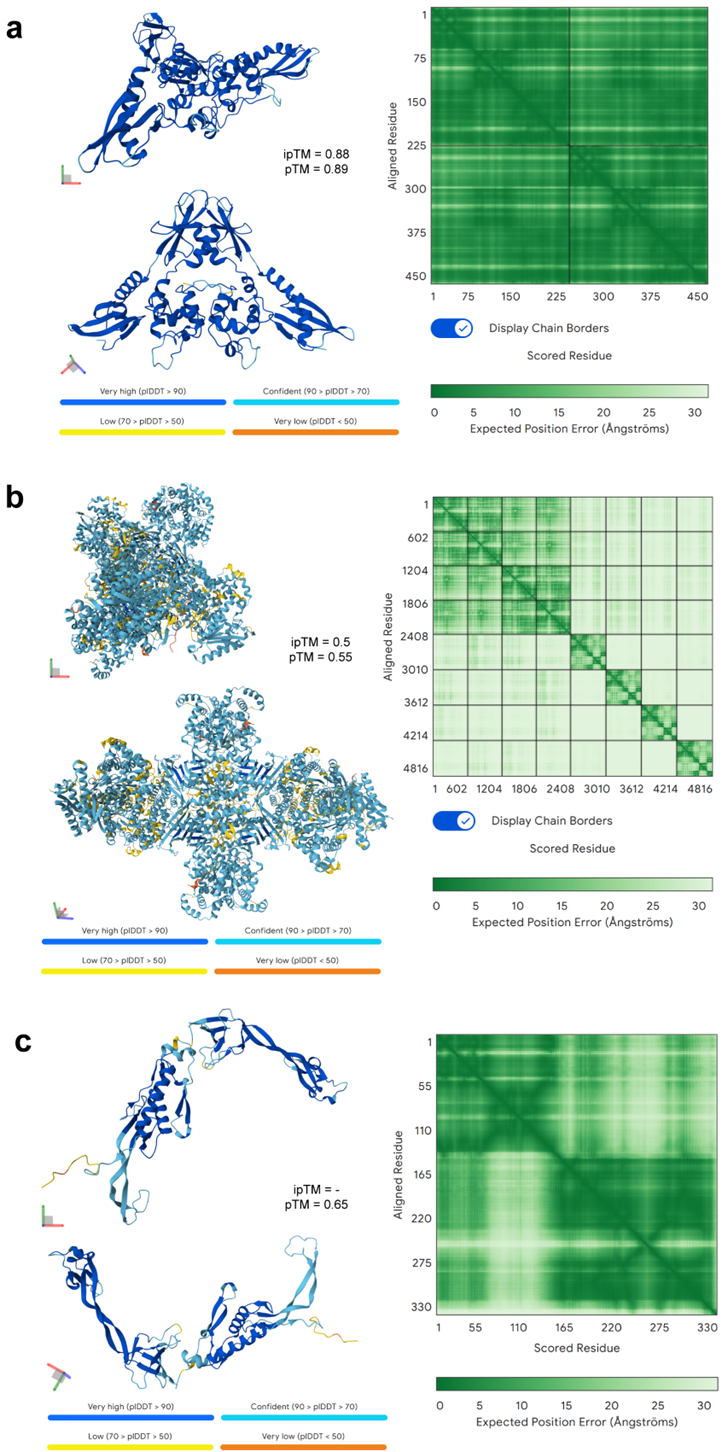

**Supplementary Figure 23.** **AlphaFold3-predicted structures and confidence scores of AcrIF24, GajAB, and Gad1.** Predicted structures of the AcrIF24 dimer **(a)**, GajAB octamer **(b)**, and Gad1 protein **(c)**, each shown from two viewing angles colored by pLDDT scores, together with the corresponding Predicted Aligned Error (PAE) plots.
